# Microbiome-based correction for random errors in nutrient profiles derived from self-reported dietary assessments

**DOI:** 10.1101/2023.11.21.568102

**Authors:** Tong Wang, Yuanqing Fu, Menglei Shuai, Ju-Sheng Zheng, Lu Zhu, Andrew T. Chan, Qi Sun, Frank B. Hu, Scott T. Weiss, Yang-Yu Liu

## Abstract

Since dietary intake is challenging to directly measure in large-scale cohort studies, we often rely on self-reported instruments (e.g., food frequency questionnaires, 24-hour recalls, and diet records) developed in nutritional epidemiology. Those self-reported instruments are prone to measurement errors, which can lead to inaccuracies in the calculation of nutrient profiles. Currently, few computational methods exist to address this problem. In the present study, we introduce a deep-learning approach --- **<u>M</u>**icrobiom**<u>e</u>**-based nu**<u>t</u>**rient p**<u>r</u>**of**<u>i</u>**le **<u>c</u>**orrector (METRIC), which leverages gut microbial compositions to correct random errors in self-reported dietary assessments using 24-hour recalls or diet records. We demonstrate the excellent performance of METRIC in minimizing the simulated random errors, particularly for nutrients metabolized by gut bacteria in both synthetic and three real-world datasets. Further research is warranted to examine the utility of METRIC to correct actual measurement errors in self-reported dietary assessment instruments.

## Introduction

An unhealthy diet can increase the risk of many diseases^1,2,3,4^. For instance, excess intake of sugar or saturated fat could elevate the risk of coronary heart disease^5–8^. The investigation of the association between poor dietary habits and chronic diseases requires an accurate assessment of dietary intake in large population samples. Typically, epidemiologic studies rely on self-reported instruments such as food frequency questionnaires (FFQ)^9^, Automated Self-Administered 24-hour Dietary Assessment Tool (ASA24)^10^, and 7-Day Dietary Record (7DDR)^11^, to collect diet data in large populations. However, self-reported instruments are subject to both random and systematic measurement errors^12–15^. While random errors are largely due to day-to-day individual variations in dietary intakes, systematic errors can be caused by under- or over-reporting and inaccurate estimation of portion sizes^16^. Measurement errors in the assessed food intake will be naturally propagated forward to the computation of the nutrient profile, causing inaccuracies in the assessed nutrient profile (Fig. 1a). Correcting such errors in the nutrient profile is crucial for improving the quality of nutritional epidemiology research. Several methods, such as regression calibrations^17,18^ and cumulative averages^19^ using repeated dietary assessments, have been used to correct for random errors of habitual diets in nutritional studies. These methods involve using a regression model to map the habitual dietary intake measured by a less accurate method (such as FFQ) to a more accurate habitual dietary measurement method (such as the average of 7DDRs). However, it is important to note that these methods are designed specifically to address measurement errors in habitual dietary intake and are incapable of correcting random measurement errors in single-day dietary assessments.

**Figure 1:**
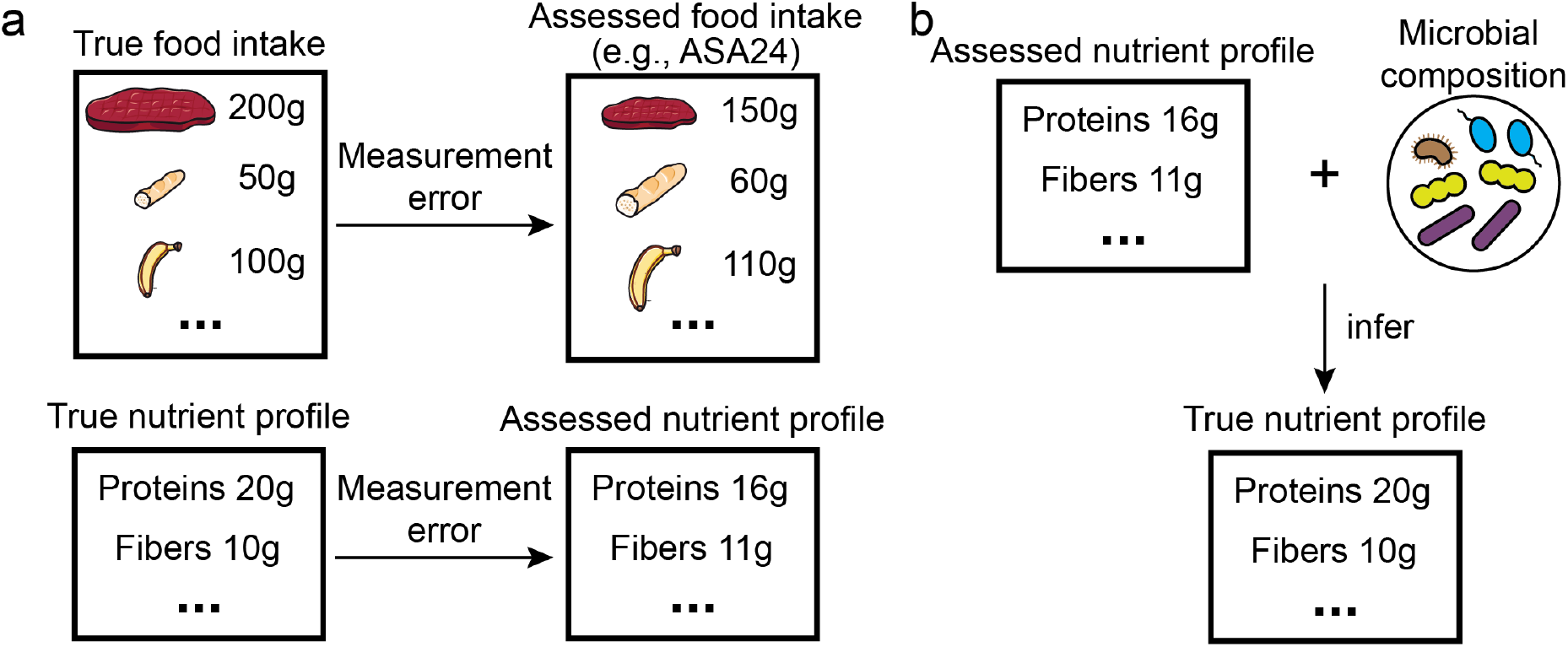
The idea of inferring the true nutrient profile based on the assessed nutrient profile and microbial composition. **a**, The typical dietary assessment such as ASA24 (Automated Self-Administered 24-hour Dietary Assessment Tool) often induces a measurement error in the food intake due to unreliable mental recall. Such a random measurement error or systematic error/bias in the food intake is carried to the nutrient profile when the food profile is converted to the nutrient profile. **b**, Our goal in this study is to remove the random part of measurement error. More specifically, we would like to infer true nutrient profiles based on assessed nutrient profiles and microbial compositions.

Signal reconstruction from corrupted or incomplete measurements is a common problem in various fields of research. For example, image denoising (i.e., removing noise from a noisy image to restore the true image) is an important topic in computer vision. The recent advance in deep neural networks has led to new methods that learn to map corrupted images to unobserved clean ones^20,21^. However, training those methods typically requires clean images, which are difficult to obtain in many cases. To overcome this challenge, a new class of methods that only leverages noisy images in training has been developed^22,23^. One noticeable example is Noise2Noise^22^, which restores clean images by only being trained on corrupted ones. When the noise is random with a mean of zero, its performance is comparable and sometimes even better than other methods trained using clean images^22^. The Noise2Noise’s underlying principle involves training the model on pairs of noisy images as both input and output. This strategy forces the neural network to predict the average of the corrupted images, thereby statistically converging the prediction of Noise2Noise towards the clean image due to the zero-mean nature of the noise^22^.

Inspired by the success of Noise2Noise^22^, we hypothesize that we can correct the random error in the assessed nutrient profile derived from the self-reported dietary assessment without using clean data (i.e., the ground truth dietary intake). Note that we focus on correcting the nutrient profile instead of the food profile (or the original dietary assessment) because the prevalence of zero values in the food profile (i.e., no consumption for many food items) renders a big challenge for machine learning tasks, while the derived nutrient profile usually has non-zero values. Also, we focus on the correction for random errors with zero means instead of systematic bias/errors with non-zero means because effectively correcting the latter requires the ground truth dietary intake, which is typically not available.

Our key idea is to incorporate gut microbial compositions as part of the input of a deep-learning model and to infer the true nutrient profile based on the assessed nutrient profile and the measured gut microbial composition (Fig. 1b). This idea is based on the knowledge that many dietary constituents reaching the large intestine fuel the growth of gut microbes^24–26^. One example is the growth of gut microbes enabled by anaerobic fermentation of indigestible polysaccharides^24,25^. As a result, the gut microbial composition is linked to dietary intake. Advances in sequencing technologies have made it possible to determine the gut microbial composition quickly and accurately. It has been shown that fecal bacteria and metabolites can be used as biomarkers for predicting whether a few food items are introduced as dietary interventions (e.g., avocado) for healthy adults in dietary intervention studies^27,28^. Hence, using objective microbial biomarkers to infer dietary intake might complement the self-reported dietary assessment and thus reduce measurement error in dietary intake. In this work, we developed a deep-learning method: **<u>M</u>**icrobiom**<u>e</u>**-based nu**<u>t</u>**rient p**<u>r</u>**of**<u>i</u>**le **<u>c</u>**orrector (METRIC). We demonstrated the effectiveness of METRIC in removing randomly added noise to both synthetic and real data.

## Results

### Overview of METRIC

We aim to infer the true nutrient profile based on the assessed one and the measured gut microbial composition. A naive way to do this is to train a machine-learning model with the assessed nutrient profiles and microbial compositions as the input and the true nutrient profiles as the output. However, this is not feasible because such training requires the true nutrient profiles that are not easily available. To address this issue, we developed METRIC that does not rely on the true nutrient profile during its training but still can remove random errors in the assessed nutrient profile during testing (Fig. 2). During the training, we generated the corrupted nutrient profiles by adding random noise to the assessed nutrient profiles, and then trained METRIC to remove the added noise by taking the corrupted nutrient profiles and the measured microbial compositions as its input and generating the assessed nutrient profiles as its output (Fig. 2a). We introduced the corrupted nutrient profiles to avoid METRIC copying the assessed nutrient profiles directly to the output, thus forcing METRIC to remove the added noise. Note that the training of METRIC did not use true nutrient profiles.

**Figure 2:**
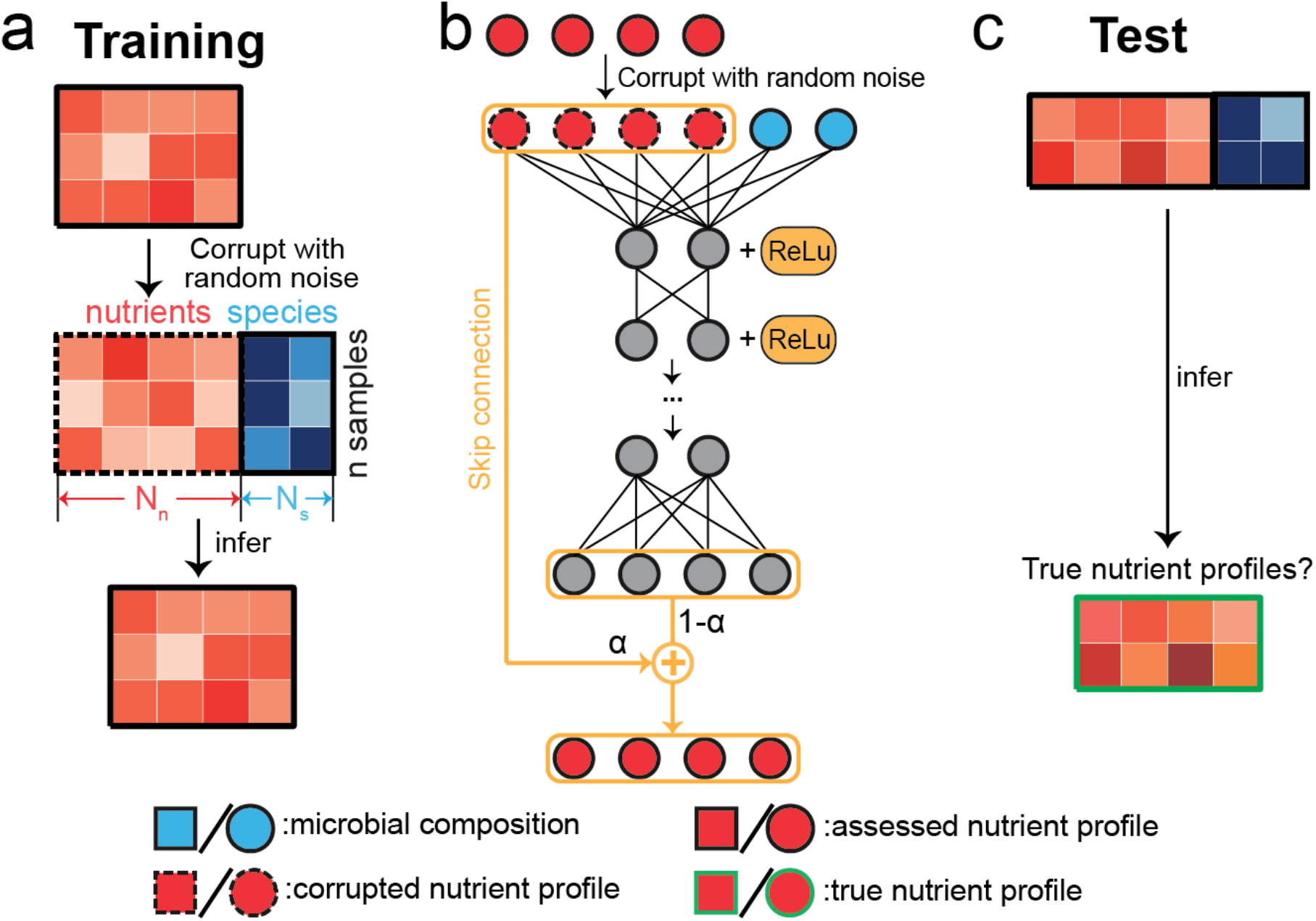
The architecture and workflow of METRIC to infer true nutrient profiles. For simplicity, we used a hypothetical example with n=3 training samples and 2 samples in the test set. For each sample, there are *N*_*s*_ microbial species and *N*_*n*_ nutrients. Across panels, microbial species and their relative abundances are colored blue. Nutrients and their amounts are colored red. The corrupted nutrient profiles are created by adding different types of random noise (i.e., Gaussian, Uniform, etc.) to the assessed nutrient profiles. Icons associated with assessed/corrupted nutrient profiles are bounded by solid black/dashed lines. Icons associated with true nutrient profiles are bounded by solid green lines. **a**, During the training of METRIC, the method takes corrupted nutrient profiles and microbial compositions as the input and learns to infer assessed nutrient profiles. **b**, Similar to multilayer perceptrons, METRIC has several hidden layers in the middle. The skip connection provides the corrupted nutrient profile directly to one layer before the final output, enabling it to skip the propagation through the hidden layers. The skipped corrupted nutrient profile multiplied by the weight parameter *α* and the final hidden layer (the bottom grey nodes) multiplied by (1 − *α*) add up as the final output (the bottom red nodes). **c**, The well-trained METRIC is applied to the test set to generate predictions for nutrient profiles whose values are compared to true nutrient profiles.

The architecture of METRIC is a neural network that consists of three hidden layers, in addition to its input and output layers (Fig. 2b). Each hidden layer has a fixed dimension of 256. The link weights in the neural network are initialized using the Xavier Initialization. The training loss is the mean squared error. The predictive performance is assessed by the mean Pearson correlation coefficient between true and predicted nutrient concentrations averaged across all nutrients 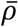. One unique feature is the skip connection that adds the corrupted nutrient profiles directly to the final output of neural networks, which previously has been shown to enhance the training and predictive performance for deep neural networks^29^. Similar to the existence of the skip connection in Noise2Noise^22^, the introduction of the skip connection enables the neural networks to adjust the prediction based on the corrupted nutrient profile, ensuring that output variables would not deviate too much from the corrupted nutrient profile. More details about the architecture can be found in the Methods section. Overall, METRIC is a generic “denoiser” that learns to remove any random noise added to the nutrient profile. With this generic ability to remove random noise, the well-trained METRIC should be able to remove random measurement errors by generating predictions closer to the true nutrient profiles when it takes the assessed nutrient profiles and microbial compositions as its input (Fig. 2c).

We split each dataset into two non-overlapping parts: a training and a test set. METRIC was trained on the training set and then used to generate predictions for the test set. The correction performance is measured by comparing the predicted nutrient profiles with the “true” nutrient profiles in the test set (how we obtain the “true” nutrient profiles for each scenario will be explained in separate sections below). To measure the predictive performance of each nutrient, we adopted the Pearson Correlation Coefficient *ρ* between its predicted and true corrected values.

### METRIC can reduce measurement errors in nutrient profiles of synthetic data

We first validated METRIC using synthetic data for which we know the ground truth. We used the Microbial Consumer-Resource Model (MiCRM)^30^ to generate three types of data: (1) *true* nutrient profiles (i.e., the ground-truth nutrient consumption), (2) *assessed* nutrient profiles (i.e., the true nutrient profiles with random noise added to mimic measurement errors), and (3) *corrupted* nutrient profiles (i.e., the assessed nutrient profiles with artificially added random noise). MiCRM simulates the process of nutrient consumption by microbes and the following microbial growth^30^. Note that the random noise added to assessed and corrupted nutrient profiles are different. For simplicity, we only considered the nutrient consumption in MiCRM and did not model the nutrient production because most dietary nutrients cannot be produced by microbes. We created different samples by randomly sampling nutrient fluxes and then running the community assembly until we achieved a steady state. We consider sampled ground-truth nutrient fluxes as true nutrient profiles and the steady-state microbial abundances as microbial compositions. Gaussian noise N(0, *σ*^2^) with the mean of zero and standard deviation *σ* is added to true nutrient profiles to create assessed nutrient profiles. More details about MiCRM, the generation of synthetic data, and added noise can be found in the Methods section.

We generated 250 samples by simulating the assembly process for 250 independent communities with 20 nutrients and 20 bacterial species. We trained METRIC on 200 samples and tested it on the remaining 50 samples. We measured how assessed values and corrected values (i.e., predicted values on the test set) respectively correlate with true values. For each nutrient, we measured ρ between its corrected values from predictions and its true concentration (denoted as ρ_c_). Similarly, we calculated ρ between its assessed concentrations and its true concentration (denoted as ρ_a_). As the standard deviation *σ* of the Gaussian noise increases, METRIC starts to correct the introduced noise, represented by the switch from negative values of (ρ_c_− ρ_a_) to positive values in Fig. 3a. It is natural to expect that the correction of nutrient profiles with weaker noises does not work (ρ_c_< ρ_a_ for small *σ* in Fig. 3a), because the correction is not necessary when the measurement error is small (e.g., when ρ_a_ > 0.8). We focus on the case of *σ* = 1.5 from now on. The difference between the two metrics (i.e., ρ_c_− ρ_a_) reflects the correction performance. The nutrient in Fig. 3d-e is slightly corrected (ρ_c_− ρ_a_ = 0.01), while the nutrient in Fig. 3f-g is strongly corrected (ρ_c_− ρ_a_ = 0.09). Most nutrients have a better alignment between their corrected values and true values (Fig. 3h).

**Figure 3:**
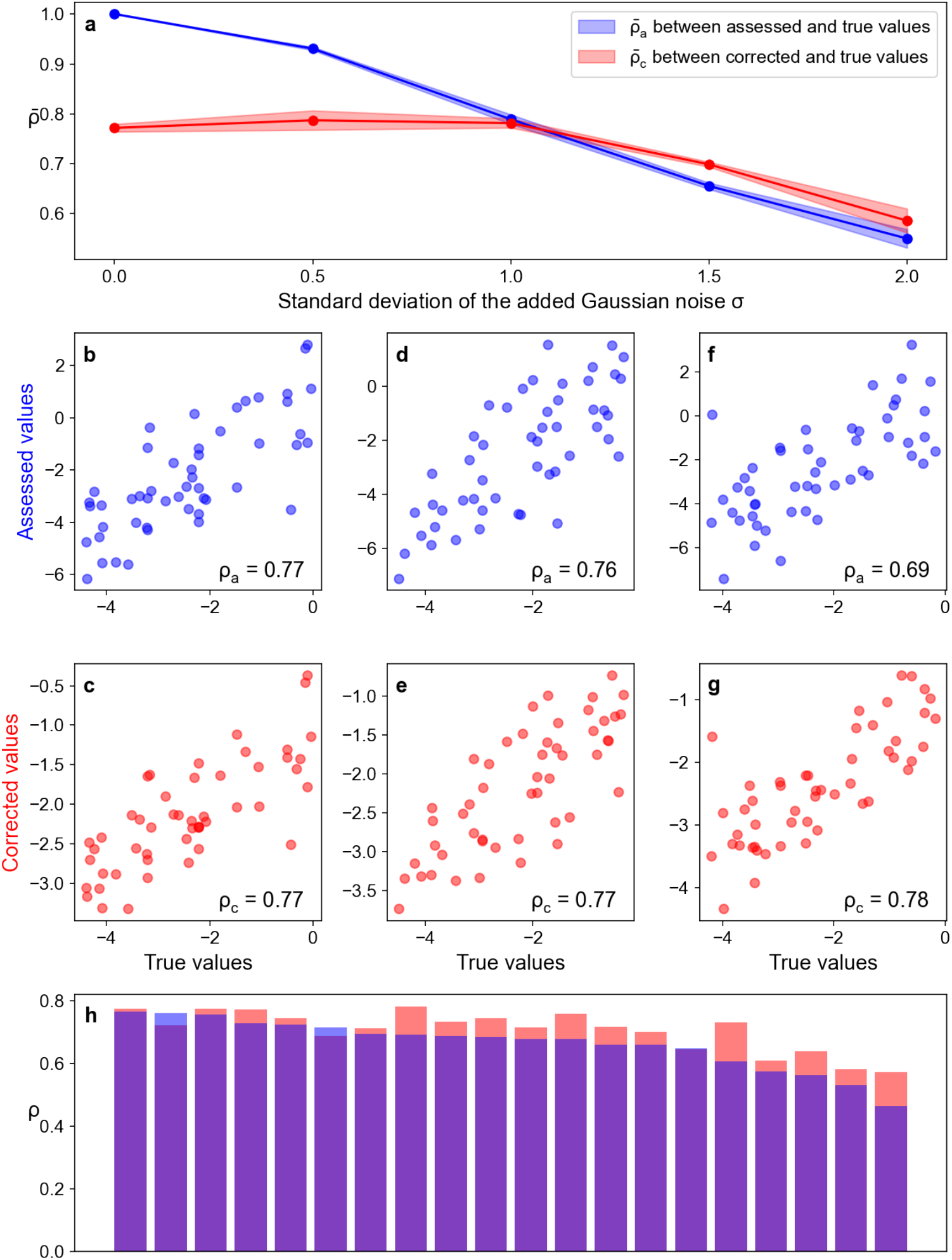
METRIC can correct the measurement error in assessed nutrient profiles on synthetic data from MiCRM^30^. The Pearson’s Rank Correlation Coefficient *ρ* is adopted to evaluate the correlation across various types of nutrient profiles. All corrected/true values shown are the log of nutrient concentrations. **a**, *ρ*_*c*_ (i.e., *ρ* between corrected and true values) and *ρ*_*a*_ (i.e., *ρ* between assessed and true values) decrease as the standard deviation of added Gaussian noise *σ* increases. All following panels focus on the case of *σ*=1.5. **b**, The correlation between assessed values and true values of log concentrations of one nutrient among different samples. **c**, The correlation between corrected values (predictions of METRIC) and true values of log concentrations of the same nutrient among different samples. Similar comparisons for another two nutrients are shown in **d-e** and **f-g. e**, The correction performance of all nutrients is measured by (*ρ*_*c*_− *ρ*_*a*_).

### METRIC mitigates measurement errors added to nutrient profiles in real data

Next, we tested METRIC on three real-world datasets. The first dataset, MCTS (MiCrobiome dieT Study), comes from a unique study that investigated the influence of diets on gut microbial composition^31^. It is unique because a large number of samples (n=210) of paired nutrient profiles and microbial compositions were collected. The nutrient profiles were calculated from ASA24 (see Methods). Different from the availability of true nutrient profiles in synthetic data, the true nutrient profiles are not available for real data. To deal with this issue, we treated the nutrient profiles derived from ASA24 as the “true” nutrient profiles and added random noise (Gaussian noise with the mean of zero and standard deviations *σ*) to them as “assessed” nutrient profiles. As *σ* increases, METRIC starts to better correct the introduced noise (Fig. 4a). We focus on the case of *σ* = 1.0 from now on. We found that carotene has a large *ρ*_a_ and is not improved by METRIC (*ρ*_a_ = 0.99 versus *ρ*_c_ = 0.97 in Figs. 4b-c). By contrast, fiber has a small ρ_a_ and is strongly improved (ρ_a_ = 0.35 versus ρ_c_= 0.58 for Figs. 4f-g). We believe that the large correction in the total fiber intake was due to most fibers being digested by gut microbes^24,25^. Overall, nutrients with smaller ρ_a_ have better correction performance, and nutrients with large ρa rarely improved (Fig. 4h). The mean correction performance averaged over all nutrients is 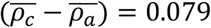.

**Figure 4:**
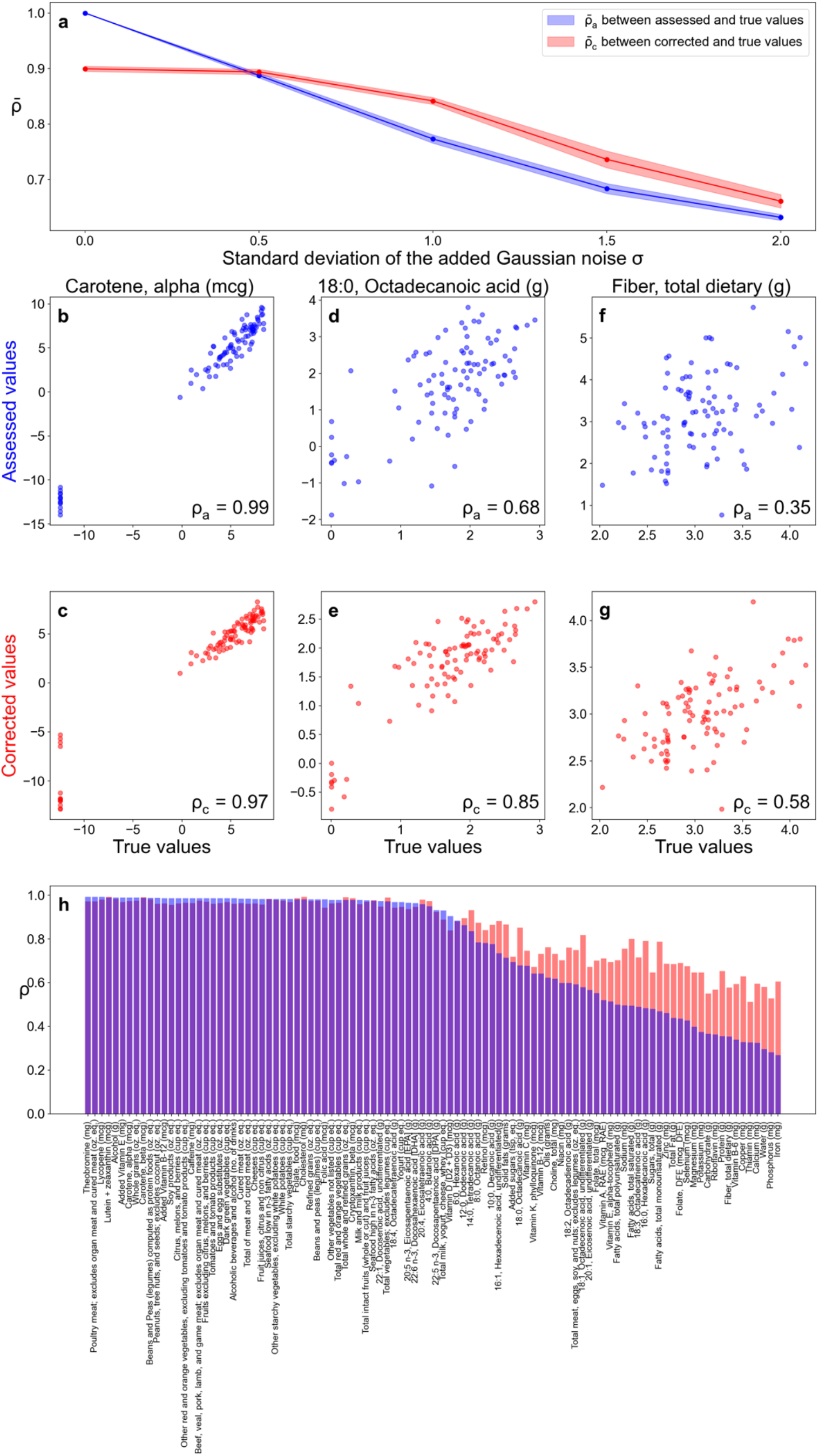
METRIC can correct the measurement error in assessed nutrient profiles on real data from MCTS^31^. The Pearson’s Rank Correlation Coefficient *ρ* is adopted to evaluate the correlation across various types of nutrient profiles. All nutrient concentrations are in the unit of grams. All corrected/true values shown are the log of nutrient concentrations. **a**, *ρ*_*c*_ (i.e., *ρ* between corrected and true values) and *ρ*_*a*_ (i.e., *ρ* between assessed and true values) decrease as the standard deviation of added Gaussian noise *σ* increases. All following panels focus on the case of *σ*=1.0. **b**, The correlation between assessed values and true values of log concentrations of carotene among different samples. **c**, The correlation between corrected values (predictions of METRIC) and true values of log concentrations of carotene among different samples. **d-e**, The similar comparison for octadecanoic acid shows a modest correction. **f-g**, The similar comparison for fiber shows a strong correction. **h**, The correction performance for all nutrients is measured by (*ρ*_*c*_ − *ρ*_*a*_).

We also tried to run METRIC without using the microbial composition, finding that the correction performance 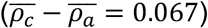 is worse than that with included microbial composition (Supplementary Fig. 1). We also computed the sensitivity which is defined as the ratio between the reduction in *ρ* of nutrient *α* and the perturbation amount of species *i* (Supplementary Fig. 2), which could detect some possible interactions between nutrients and species. For example, the sensitivity of monounsaturated fatty acids towards *Bacteroides uniformis* is large (∼0.025, larger than 99.98% of the inferred sensitivity values), and is supported by the previously observed reduction of monounsaturated fatty acids by *Bacteroides uniformis*^32^. In addition, our sensitivity analysis revealed that the top three microbial taxa linked to fiber content correction, which exhibit the highest sensitivity values, are well-documented as fiber degraders in the literature: *Bacteroides plebeius*^33^, *Parabacteroides sp*^34^, and *Bacteroides sp*^35^.

Then, we applied METRIC to the second dataset MLVS (Men’s Lifestyle Validation Study)^36,37^. Specifically, we utilized the composition of gut microbiomes and the one-day dietary assessment of the 7-day dietary records (7DDRs). The 7DDRs are widely recognized to be the most reliable estimation of dietary intake because participants are required to measure and report gram weights for foods both before they start eating and after they finish, thereby enabling the calculation of the actual food consumption based on the difference in weight^38^. To guarantee the usefulness of gut microbial composition in correcting dietary assessment, we required the identification of paired microbial compositions and dietary assessments with matching dates. In MLVS, a total of 599 paired samples with matching dates were found. Similarly, as we did for the MCTS dataset, we regarded the nutrient profile derived from the 7DDRs as the “true” nutrient profile and added varying levels of Gaussian noise to it as the “assessed” nutrient profile. Then we trained METRIC on 80% of the data and tested it on the remaining 20%. Consistent with our previous findings, the trained METRIC exhibits an ability to correct the nutrient profile (Fig. 5a), especially for large *σ*. For the case of *σ* = 1.0, the mean correction performance 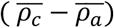 is 0.072 (Fig. 5h). Across all nutrients, dietary fiber was the strongest corrected nutrient (Figs. 5f-g).

**Figure 5:**
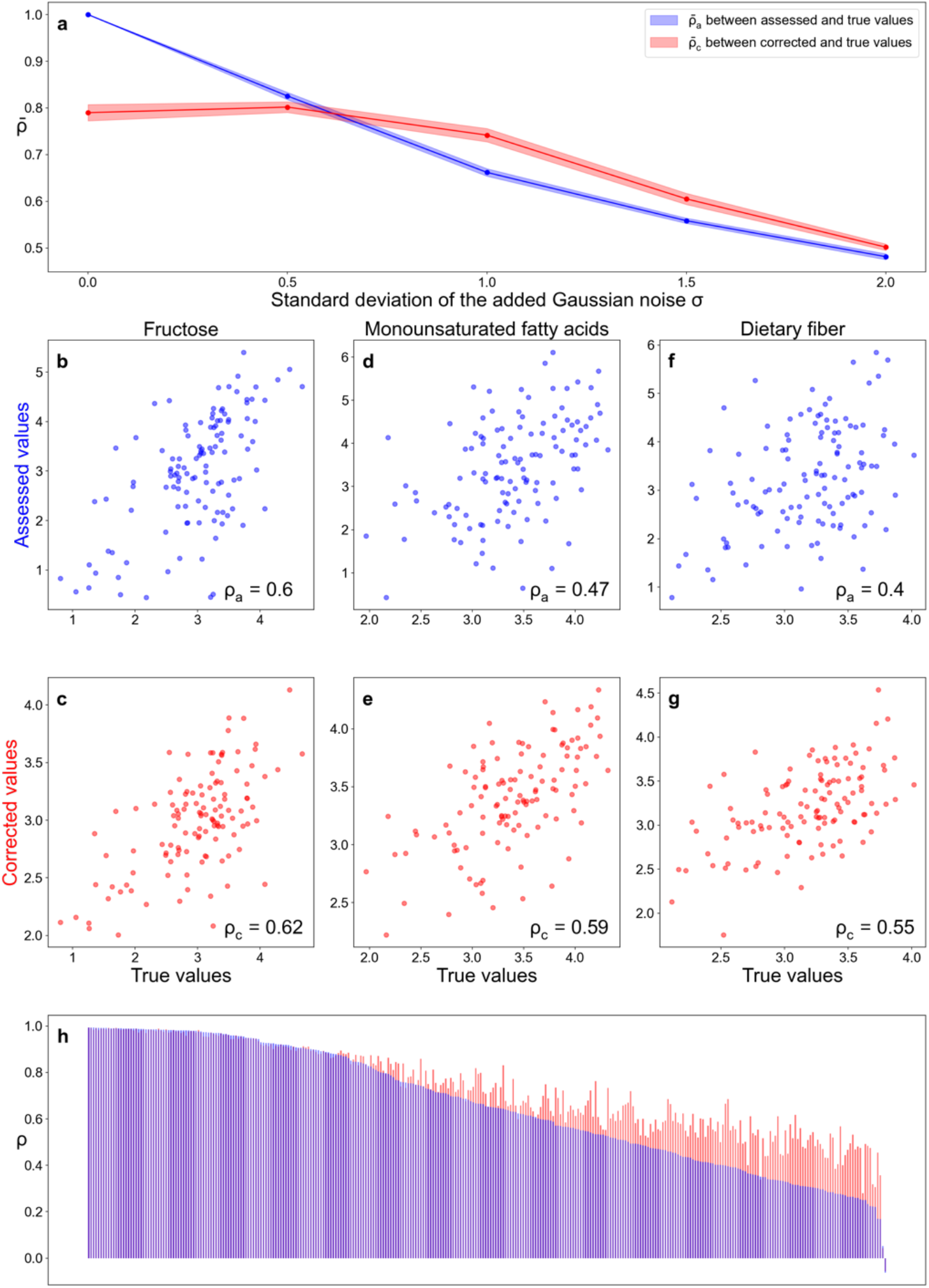
METRIC can correct the measurement error in assessed nutrient profiles from MLVS^36,37^. The Pearson’s Rank Correlation Coefficient *ρ* is adopted to evaluate the correlation across various types of nutrient profiles. All nutrient concentrations are in the unit of grams. All corrected/true values shown are the log of nutrient concentrations. **a**, *ρ*_*c*_ (i.e., *ρ* between corrected and true values) and *ρ*_*a*_ (i.e., *ρ* between assessed and true values) decrease as the standard deviation of added Gaussian noise *σ* increases. All following panels focus on the case of *σ*=1.0. **b**, The correlation between assessed values and true values of log concentrations of fructose among different samples. **c**, The correlation between corrected values (predictions of METRIC) and true values of log concentrations of fructose among different samples. **d-e**, The similar comparison for monounsaturated fatty acids shows a modest correction. **f-g**, The similar comparison for dietary fiber shows a strong correction. **h**, The correction performance for all nutrients is measured by (*ρ*_*c*_ − *ρ*_*a*_). Nutrient names are not added due to lack of space.

Finally, we applied METRIC to the third dataset WE-MACNUTR (Westlake N-of-1 Trials for Macronutrient Intake)^39^. WE-MACNUTR is a dietary intervention study that implemented a ‘complete feeding’ strategy, providing three isocaloric meals per day to 28 participants over a span of 72 days. Each participant completed high-fat, low-carbohydrate and low-fat, high-carbohydrate diets in a randomized sequence, with a 6-day wash-out period between them. Since the diets were completely controlled and well-documented, the nutrient profile derived from this dataset closely reflects the true nutrient profile. In WE-MACNUTR, we found 317 paired samples with both microbial compositions and nutrient profiles. Considering the nutrient profile from the complete feeding as the true nutrient profile, we introduced varying levels of noise (Gaussian noise N(0, *σ*^2^) with different standard deviations *σ*) to create the assessed nutrient profile. Like our earlier results, METRIC can correct the nutrient profile when *σ* is large (Fig. 6a). When *σ* = 1.0, the mean correction performance 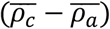 is 0.118 (Fig. 6h) and dietary fiber again exhibits a substantial correction (Figs. 6f-g).

**Figure 6:**
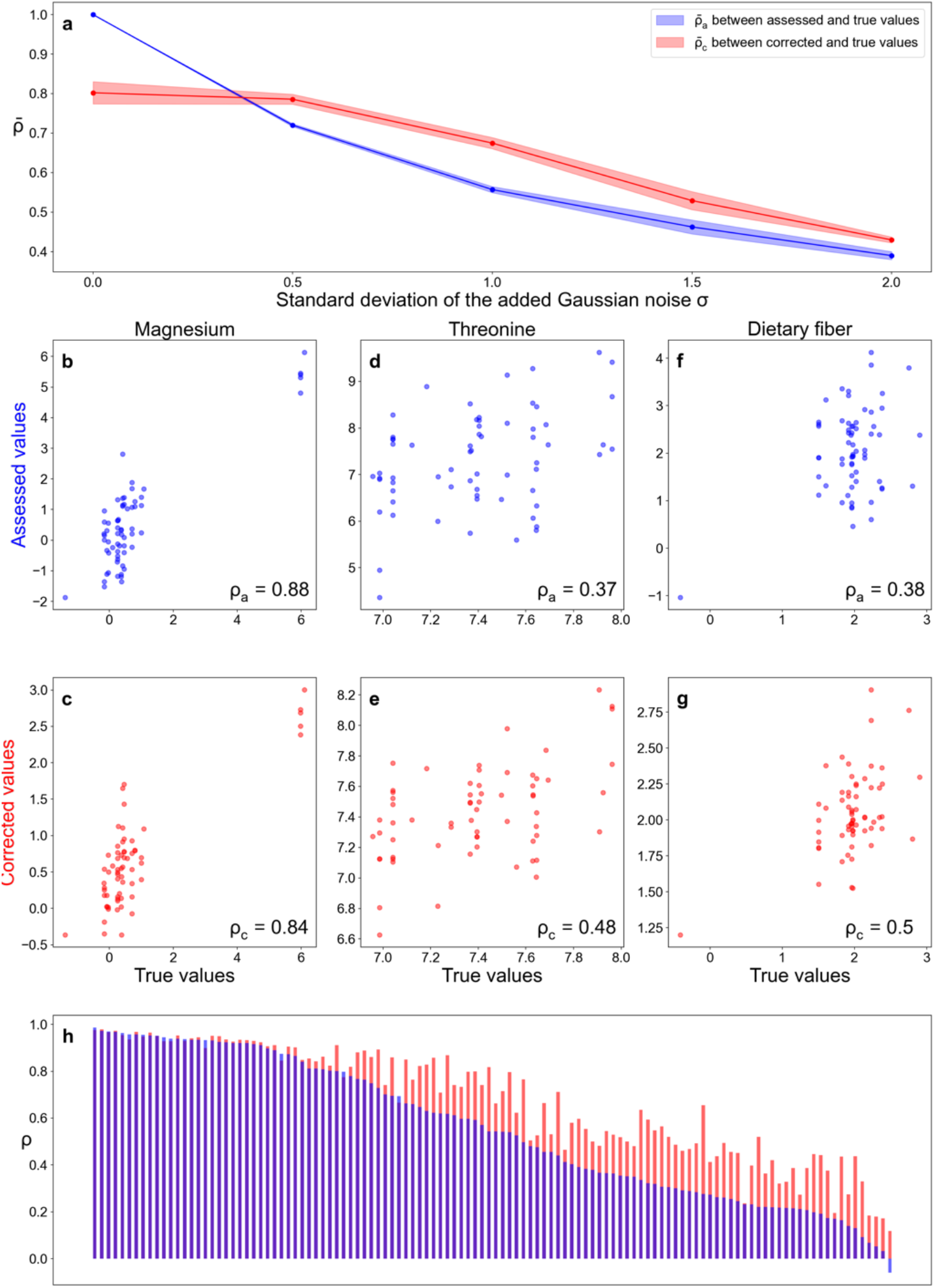
METRIC can correct the measurement error in assessed nutrient profiles from WE-MACNUTR^39^. The Pearson’s Rank Correlation Coefficient *ρ* is adopted to evaluate the correlation across various types of nutrient profiles. **a**, *ρ*_*c*_ (i.e., *ρ* between corrected and true values) and *ρ*_*a*_ (i.e., *ρ* between assessed and true values) decrease as the standard deviation of added Gaussian noise *σ* increases. All nutrient concentrations are in the unit of grams. All corrected/true values shown are the log of nutrient concentrations. All following panels focus on the case of *σ*=1.0. **b**, The correlation between assessed values and true values of log concentrations of magnesium among different samples. **c**, The correlation between corrected values (predictions of METRIC) and true values of log concentrations of magnesium among different samples. **d-e**, The similar comparison for threonine shows a modest correction. **f-g**, The similar comparison for dietary fiber shows a strong correction. **h**, The correction performance for all nutrients is measured by (*ρ*_*c*_ − *ρ*_*a*_). Nutrient names are not added due to lack of space.

To provide a more representative picture of METRIC’s robustness and effectiveness across different noise levels, we also investigated situations where the correction is less effective for all three datasets (standard deviation of the noise *σ* = 0.5; Supplementary Figs. 3-5). Although the overall correction performance 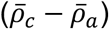 is weak when *σ* = 0.5, the correction still works well for nutrients with smaller *ρ*_*a*_ (e.g., dietary fibers). We also evaluated the predictive performance using a more quantitative metric, the mean absolute error. We found that the overall pattern in correction performance measured by mean absolute error aligns with that measured by Pearson correlation across datasets (Supplementary Figs. 6-8). For the MLVS and WE-MACNUTR datasets, we also tried to evaluate METRIC’s correction performance without using the microbial composition, finding that the correction performance is comparable to that achieved when the microbial composition is included (Supplementary Figs. 9&10). This implies that METRIC can still be leveraged to correct nutrient profiles even in the absence of gut microbial compositions.

We capitalized on the longitudinal nature of the MCTS dataset to explore whether increasing temporal offsets between microbiome and diet data impacts the correction efficiency of our method. Specifically, we increased the offset by aligning the diet of day *t* with the microbiome of day *t* + Δ*t* and subsequently correcting nutrient profiles. Our analysis reveals that the correction performance progressively decreases as the offset Δ*t* deviates from 1 day (Supplementary Fig. 11). This serves as a validation of METRIC, as it indicates that microbiome-diet relationships are causal.

Given the absence of ground-truth nutrient profiles for direct validation, we instead conducted an indirect validation of our method by checking if the noise level of real-life nutrient profiles is within the regime where we can remove the random measurement errors well. We found that the correction performance of METRIC is great when the mean Pearson correlation coefficient 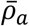 a is below 0.8 (Figs. 4-6). Due to the lack of ground-truth nutrient profiles to directly quantify the noise level, we can only approximate this indirectly by reflecting the nutrient variability using the multiple-day 7DDRs in MLVS. Specifically, we calculated the Pearson correlation coefficient *ρ* between concentrations of a nutrient derived from one 7DDR and its average values obtained from multiple 7DDRs for seven consecutive days (Supplementary Fig. 12a). We found that the mean Pearson correlation coefficient 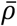 is 0.77, which is below 0.8. Additionally, 62.0% of 329 nutrients have *ρ*< 0.8. A similar analysis on the dataset MCTS revealed that 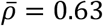 and 95.0% of nutrients have *ρ*< 0.8 (Supplementary Fig. 12b). The WE-MACNUTR dataset was not analyzed due to the absence of dietary assessments. These findings across both datasets confirm that the approximated noise levels are within a range where METRIC is effective at correcting random measurement errors.

We also examined the impact of noise with a non-zero mean by introducing the Gaussian noise N(*μ, σ*^2^). For the three datasets we used, we set *σ* = 1 as this is the regime where our method, METRIC, consistently shows strong correction performance. We then gradually increased the mean of the noise *μ* from 0.0 to 2.0 to create the assessed nutrient profile. When applying METRIC to remove the noise, we observed that its correction performance diminishes to zero as *μ* increases (Supplementary Figs. 13-15), indicating that METRIC can remove the random error but not the systematic drift or shift.

## Discussion

We presented a deep-learning method, METRIC, to correct random simulated measurement errors in the nutrient profile. The method relies on the assessed nutrient profile and microbial composition to learn how to infer the true nutrient profile. First, we validated its performance on synthetic data where we directly modeled true and assessed nutrient profiles. Then we applied METRIC to three distinct real clinical datasets with added noise to the nutrient profile and found that it can correct the nutrient profile well, especially for nutrients with large errors or metabolized by gut microbes. Similar to the class of computational methods that denoise images even without clean targets^22,23^, METRIC offers a significant advantage by being capable of correcting the random measurement error without using the true nutrient profiles during the training. This attribute makes our method particularly useful in real-life scenarios where only assessed nutrient profiles are available, without access to true nutrient profiles.

We admit that METRIC has several limitations. First, without using clean targets (i.e., the ground truth dietary intake), METRIC is only capable of removing random measurement errors that have zero means. It cannot remove noise with a non-zero mean, just like Noise2Noise^22^. Thus, METRIC cannot correct the systematic bias/drift/error (with a non-zero mean) in nutrient profiles. Effectively correcting the systematic bias requires both assessed and true nutrient profiles to discern the consistent deviation between them. However, it is very unlikely that both assessed and true nutrient profiles are available to measure the real systematic bias in real-world data. If both types of nutrient profiles are available, it remains to be investigated whether it is possible to design a regression model that predicts true nutrient profiles based on assessed nutrient profiles, thereby achieving the task of removing the systematic bias. Second, METRIC uses microbial compositions of fecal samples to correct nutrient profiles derived from ASA24 or 7DDR, which were collected exactly one day prior to fecal sample collection. We do not expect that microbial compositions of fecal samples collected from a particular time point could be used to correct nutrient profiles derived from FFQ, where subjects report how often each food item was consumed over a specified period, typically the past month or year. To clearly demonstrate this point, we leveraged the FFQ, 7DDR, and gut microbiome data from MLVS to compare the performance of using gut microbial compositions to predict the 7DDR- or FFQ-based nutrient profiles. We found that the predictability of FFQ-based nutrient profiles is much worse than that of 7DDR-based nutrient profiles (Supplementary Fig. 16). Consequently, we believe that METRIC cannot be used to correct random errors in FFQ-based nutrient profiles.

We recognize the challenges of applying a model trained on one dataset to another, particularly when biases like sequencing errors or differences in nutrient databases are present. However, training the method on one dataset and generating predictions for the other is possible if both datasets were collected and processed in the same way. For example, in the PRISM and NLIBD studies, two Inflammatory Bowel Disease cohorts—one from Boston (n=155) and an external validation cohort from the Netherlands (n=65)—were collected using the same protocols. This standardization enables accurate predictions of disease status in the NLIBD cohort by a machine learning method trained on the PRISM data^40^. Similarly, the deep-learning model mNODE trained on PRISM data showed great performance in predicting fecal metabolome based on gut microbial compositions from the NLIBD cohort^41^.

There are other methods to improve the accuracy of dietary intake measurement over traditional self-reported dietary assessments. For instance, digital documentation of meals through taking photos can be used to improve the accuracy of dietary assessment, though the validity of such technologies is yet to be established. In addition, nutritional biomarkers such as DNA barcodes for plants^42^ and metabolite biomarkers^28,43,44^ have been utilized to improve the assessment of food intake. A more accurate reflection of nutrient consumption can be obtained by using objective measurements of microbial composition or metabolomic profile, which can complement self-reported dietary assessment tools. Currently, an active research direction in the field of precision nutrition is to identify microbial and metabolite biomarkers for dietary intake. Although several studies attempted to predict the presence of food items based on fecal bacteria and metabolites^27,28^, the analyses were limited to several food items and no connection to nutrient profiles was examined. Further research is warranted to examine the utility of METRIC to correct actual measurement errors in self-reported dietary assessments.

## Methods

### Datasets

The first dataset we used comes from a study that investigated the association between diet and the gut microbiome^31^. The study has paired 24-h food records and fecal shotgun metagenomes from 34 healthy human subjects collected daily over 17 days, with 210 paired samples in total. The second dataset is from MLVS^36,37^ with 599 paired gut microbial compositions and 7DDRs. The third dataset is from WE-MACNUTR^39^, a dietary intervention study that provides three meals per day for all participants. The study involved 30 participants, with 2 participants withdrawing from the trial early and not included in the final analysis. It has 317 paired gut microbial compositions and true nutrient profiles of complete feeding with matching dates. For all machine learning tasks, the same five random 80/20 train-test splits were utilized to guarantee a fair comparison of methods.

### The generation of nutrient profiles based on dietary assessments

The generation of nutrient profiles generated from dietary assessments involves several steps and relies on comprehensive food composition databases such as USDA’s Food and Nutrient Database for Dietary Studies (FNDDS)^45^ or Harvard Food Composition Database (HFDB)^46^, etc. Each food item consumed is matched with a corresponding item in a food composition database. These databases provide detailed information on the nutrient content of food items, including macronutrients (e.g., fats, proteins, carbohydrates) and micronutrients (e.g., vitamins and minerals). By multiplying the amount of each food consumed by its nutrient content, the total intake of each nutrient is calculated to obtain the nutrient profiles. For example, the MCTS dataset utilized ASA24 to collect dietary intake data, where documented foods were assigned to nutrients based on the USDA’s FNDDS 2011-2012^45^. In the case of the MLVS dataset, nutrient intake was derived from 7DDRs using the Nutrition Data System for Research^47^, which provides a comprehensive food and nutrient database managed by the University of Minnesota Nutrition Coordinating Center. For the WE-MACNUTR dataset, which focuses on a Chinese population, nutrient profiles were derived from the foods directly provided to participants using the electronic version of China Food Composition Tables (Standard Edition)^48^.

### The generation of synthetic data

Similar to the traditional consumer-resource model^49,50^ and Microbial Consumer-Resource Model^30^, we simulated the process of nutrient consumption by microbes and the following microbial growth. For simplicity, we assumed a global pool with 20 microbial species and 20 nutrients in total. To mimic the difference in nutrient consumption profiles across individuals, we randomly assigned flux for each sample: to create a unique sample, the supply rate of each nutrient is randomly drawn from a uniform distribution 𝒰[0, 1]. We recorded the assigned supply rates of all nutrients as the true nutrient profile. The Gaussian noise with a mean of 0 and standard deviation of 0.5 is added to true nutrient profile to create assessed nutrient profiles with measurement errors. After the assignment of nutrient fluxes, we simulated the community assembly dynamics until the system reaches a steady state. The steady-state microbial relative abundances are recorded as the microbial composition for the sample. Then we repeated this procedure of generating samples 250 times, forming the synthetic dataset we used in this study. The 250 samples are split in the 80/20 ratio as the training/test set.

The overall population dynamics for the concentration of nutrient *N*_*α*_ and abundance of microbial species *M*_i_ can be written as follows:

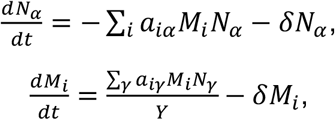

where *a*_*iα*_ is the consumption rate of nutrient *α* by the species *i, δ* is the dilution rate, and *Y* is the yield. For simplicity, we assumed *δ* = 0.1 and *Y* = 1. The *a*_*iα*_ is assumed to be non-zero with a probability of 50%. If *a*_*iα*_ is non-zero, its value is drawn from the uniform distribution 𝒰[0, 10]. Eventually, each *a*_*iα*_ is divided by the number of nutrients that can be consumed by the species *i*, avoiding the outgrowth of generalists.

### METRIC

The core of METRIC is the MLP (Multilayer Perceptron).

- Data processing: The Centered Log-Ratio transformation is applied to microbial relative abundances, and the log transformation is applied to the nutrient profiles.
- Model detail: METRIC has 3 hidden layers in the middle, sandwiched by input and output variables. Each hidden layer has a fixed hidden layer dimension of 256. The Xavier Initialization is used to initialize the weights in the neural network. A skip connection from corrupted nutrient profiles is introduced to add to the final layer of MLP. Specifically, the final prediction is the sum of (1) the corrupted input from the skip connection multiplied by a weight parameter *α* and (2) the final layer of MLP multiplied by (1 − *α*). The optimal value for *α* is chosen based on the five-fold cross-validation results on the training set.
- Training method: The Adam optimizer^51^ is used for the gradient descent. The training loss is the mean squared error. The training stops when the mean Pearson Correlation Coefficient of nutrients 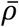 on the validation/test set starts to decrease within the past 10 epochs.
- Activation function: ReLU (Rectified Linear Unit).

### Statistics

To calculate correlation throughout the study, we used Pearson’s correlation coefficient. All simulations and analyses were performed using standard numerical and scientific computing libraries such as NumPy and SciPy in the Python programming language (version 3.7.1) and Jupyter Notebook (version 6.1).

## Data availability

We only used the data collected by existing studies. Instructions for downloading sequencing data and dietary intakes analyzed in this work can be found in the literature exploring MCTS (MiCrobiome dieT Study)^31^, MLVS (Men’s Lifestyle Validation Study)^36,37^, and WE-MACNUTR (Westlake N-of-1 Trials for Macronutrient Intake)^39^. To facilitate the data downloading, the URLs to those datasets are also provided in our GitHub repository (https://github.com/wt1005203/METRIC).

## Code availability

All code for simulations used in this manuscript can be found at https://github.com/wt1005203/METRIC.

## Acknowledgements

We acknowledge grants from the National Institutes of Health (R01AI141529, R01HD093761, R35CA253185, RF1AG067744, UH3OD023268, U19AI095219, U01HL089856, U01-152905, and U01-167552) and Cancer Grand Challenges Team PROSPECT. We thank Walter Willett, Eric Rimm, and Lorelei Mucci for valuable discussions.

## Author contributions

T.W. and Y.-Y.L. designed the project. T.W. performed all the numerical calculations and data analysis. T.W. processed the real data with assistance from Y.Fu. All authors analyzed the results. T.W. and Y.-Y.L. wrote the paper. All authors edited and approved the paper.

