## Supplemental Information for "Microbiome-based correction for random errors in nutrient profiles derived from self-reported dietary assessments"

### Supplementary Figures

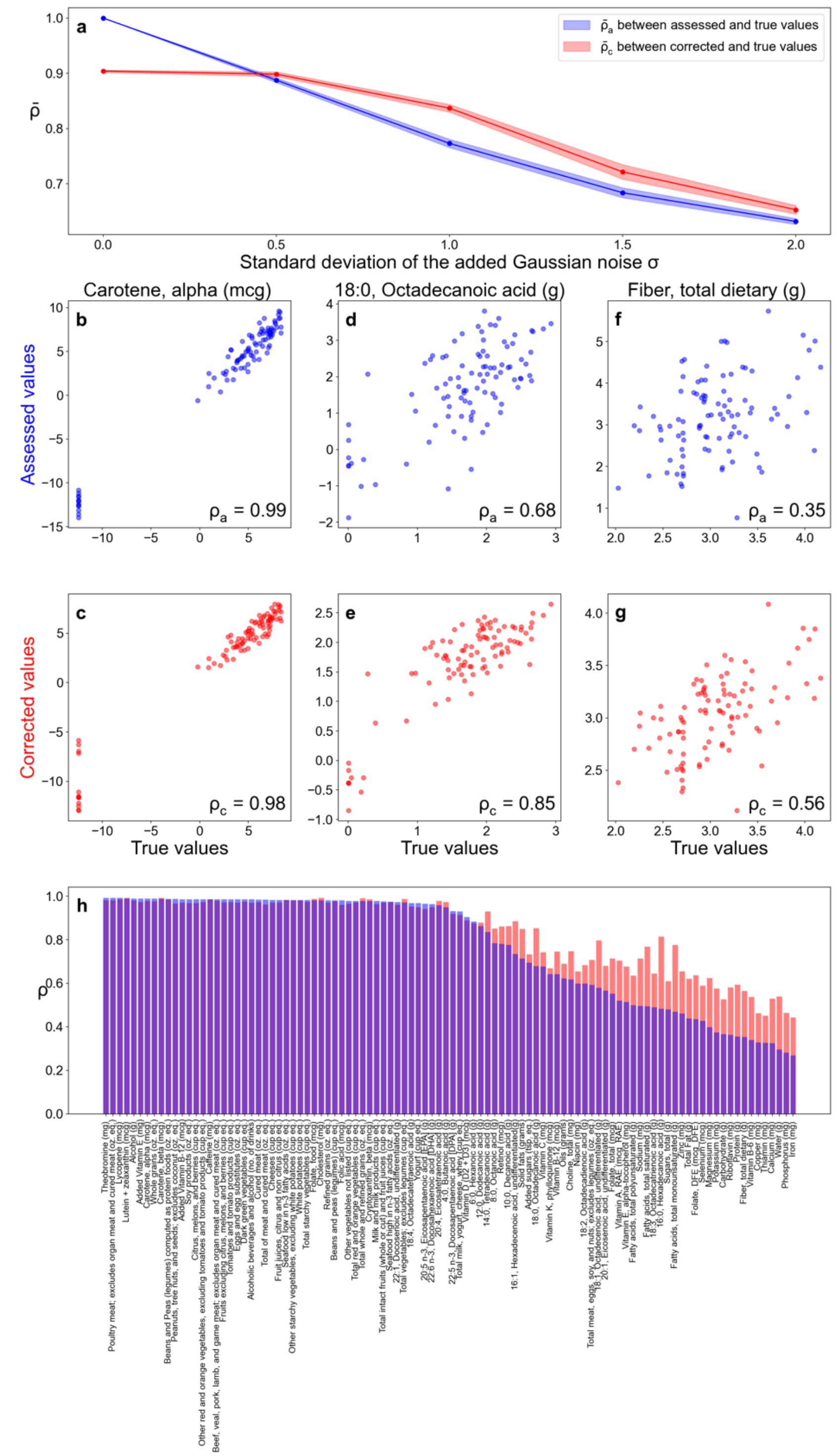


Supplementary Figure 1: **The correction performance of METRIC is weaker when the microbial composition is not included in the input on real data from MCTS[31]. The**

Pearson's Rank Correlation Coefficient $\rho$ is adopted to evaluate the correlation across various types of nutrient profiles. All nutrient concentrations are in the unit of grams. All corrected/true values shown are the log of nutrient concentrations. **a,** $\rho_c$ (i.e., $\rho$ between corrected and true values) and $\rho_a$ (i.e., $\rho$ between assessed and true values) decrease as the standard deviation of added Gaussian noise $\sigma$ increases. All following panels focus on the case of $\sigma$=1.0. **b,** The correlation between assessed values and true values of log concentrations of carotene among different samples. **c,** The correlation between corrected values (predictions of METRIC) and true values of log concentrations of carotene among different samples. **d-e,** The similar comparison for octadecanoic acid shows a modest correction. **f-g,** The similar comparison for fiber shows a strong correction. **h,** The correction performance for all nutrients is measured by $(\rho_c - \rho_a)$.

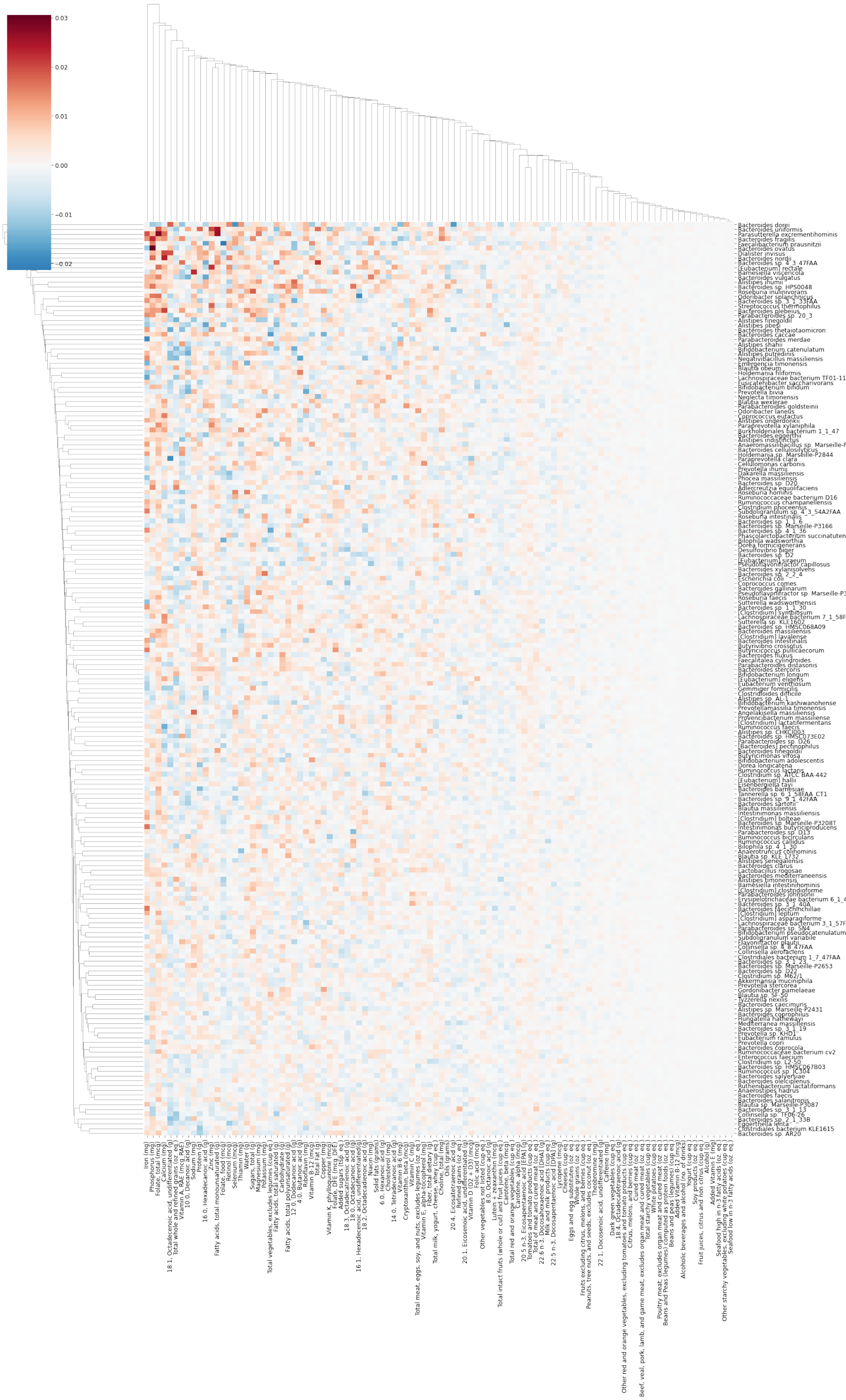
0.03
0.02
0.01
0.00
−0.01
−0.02

Supplementary Figure 2: **The sensitivity analysis reveals some important microbe-nutrient interactions on real data from MCTS[31].** The sensitivity $s_{i\beta}$ is defined as the ratio between the reduction in $\rho$ of nutrient $\beta$ and the perturbation amount of species $i$.

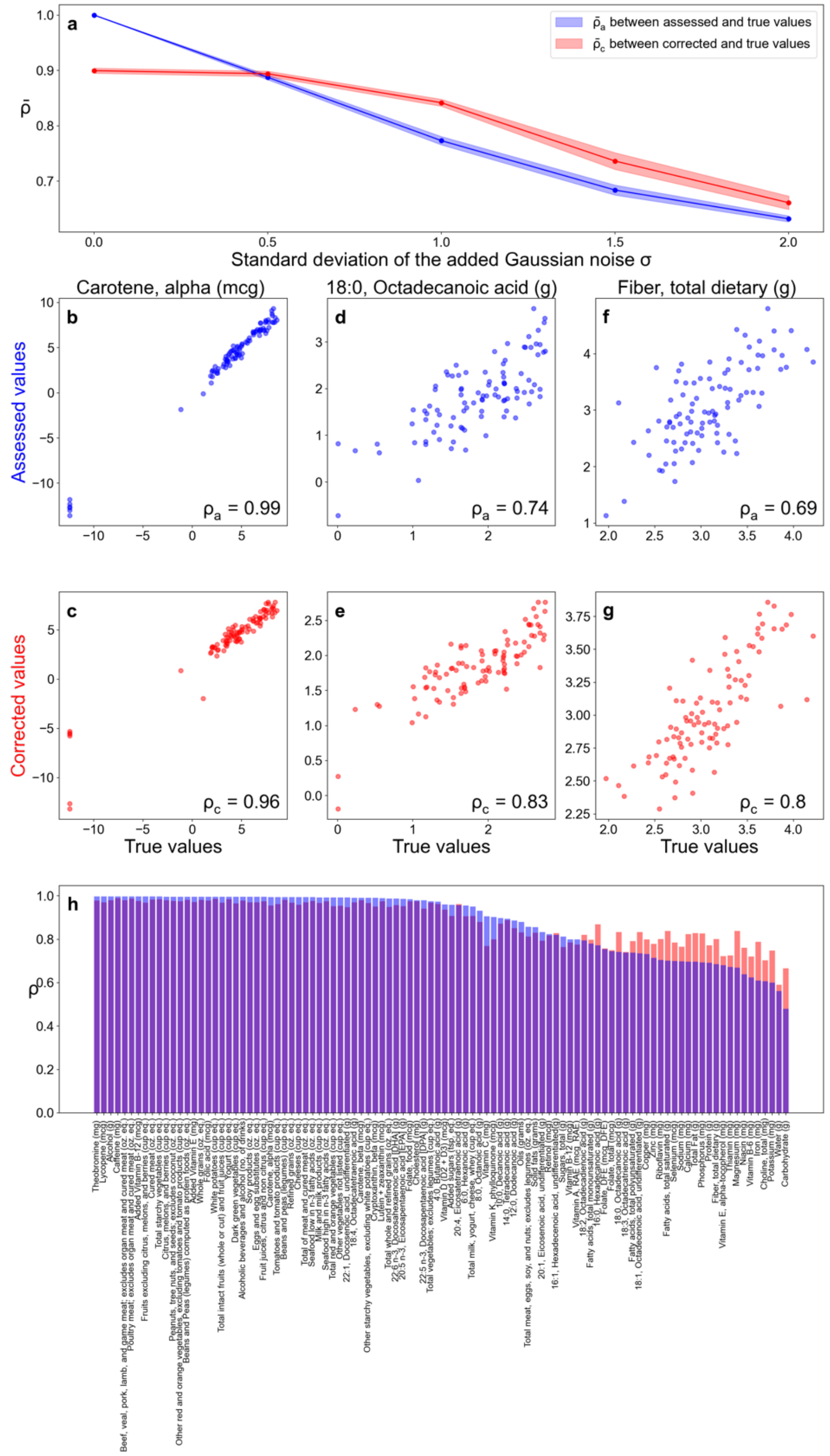


Supplementary Figure 3: **METRIC remains effective in correcting measurement errors in nutrients with large random errors, despite the overall poor correction performance when $\sigma = 0.5$ observed in the dataset MCTS[31].** The Pearson's Rank Correlation Coefficient $\rho$ is adopted to evaluate the correlation across various types of nutrient profiles. All nutrient concentrations are in the unit of grams. All corrected/true values shown are the log of nutrient concentrations. **a,** $\rho_c$ (i.e., $\rho$ between corrected and true values) and $\rho_a$ (i.e., $\rho$ between assessed and true values) decrease as the standard deviation of added Gaussian noise $\sigma$ increases. All following panels focus on the case of $\sigma$=0.5. **b,** The correlation between assessed values and true values of log concentrations of carotene among different samples. **c,** The correlation between corrected values (predictions of METRIC) and true values of log concentrations of carotene among different samples. **d-e,** The similar comparison for octadecanoic acid shows a great correction. **f-g,** The similar comparison for fiber shows a strong correction. **h,** The correction performance for all nutrients is measured by $(\rho_c - \rho_a)$.

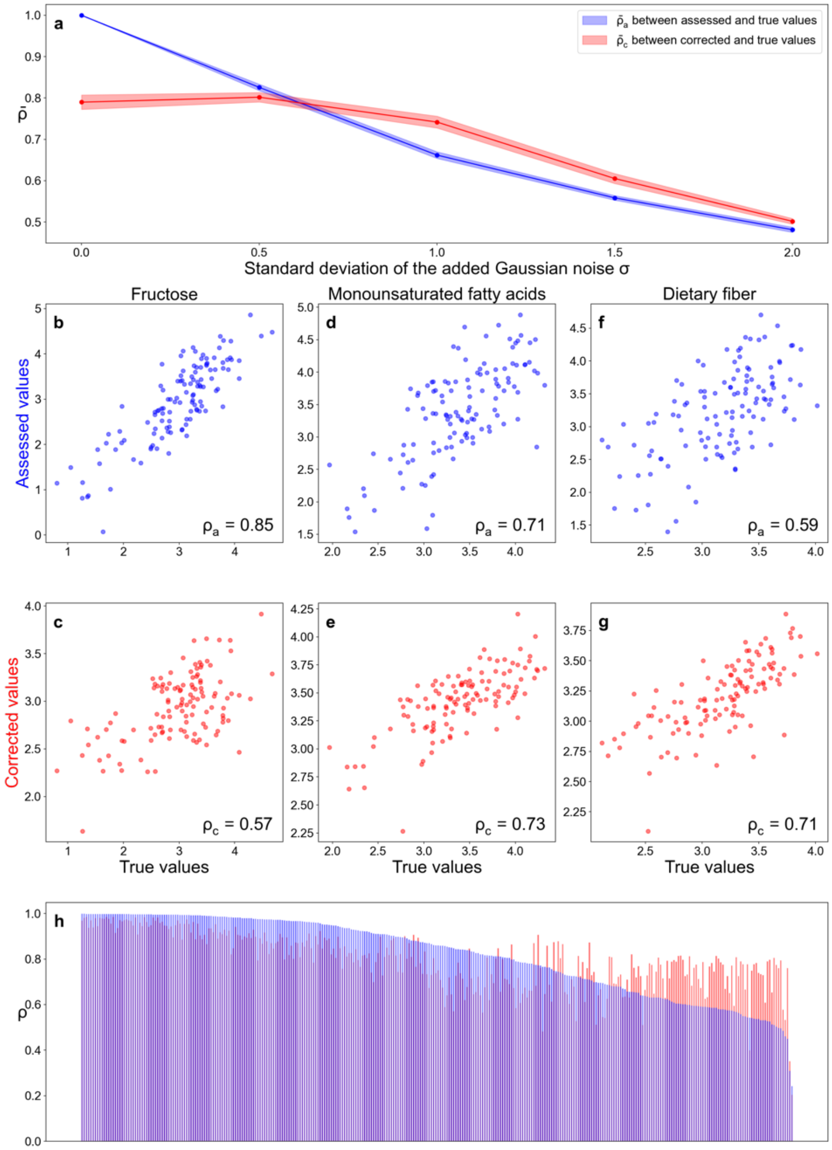


Supplementary Figure 4: **METRIC remains effective in correcting measurement errors in nutrients with large random errors, despite the overall poor correction performance when $\sigma = 0.5$ observed in the dataset MLVS**[36,37]**.** The Pearson's Rank Correlation Coefficient $\rho$ is adopted to evaluate the correlation across various types of nutrient profiles. All nutrient concentrations are in the unit of grams. All corrected/true values shown are the log of nutrient concentrations. **a,** $\rho_c$ (i.e., $\rho$ between corrected and true values) and $\rho_a$ (i.e., $\rho$ between assessed and true values) decrease as the standard deviation of added Gaussian noise $\sigma$ increases. All following panels focus on the case of $\sigma$=0.5. **b,** The correlation between assessed values and true values of log concentrations of fructose among different samples. **c,** The correlation between corrected values (predictions of METRIC) and true values of log concentrations of fructose among different samples. **d-e,** The similar comparison for monounsaturated fatty acids shows a weak correction. **f-g,** The similar comparison for dietary fiber shows a strong correction. **h,** The correction performance for all nutrients is measured by $(\rho_c - \rho_a)$. Nutrient names are not added due to lack of space.

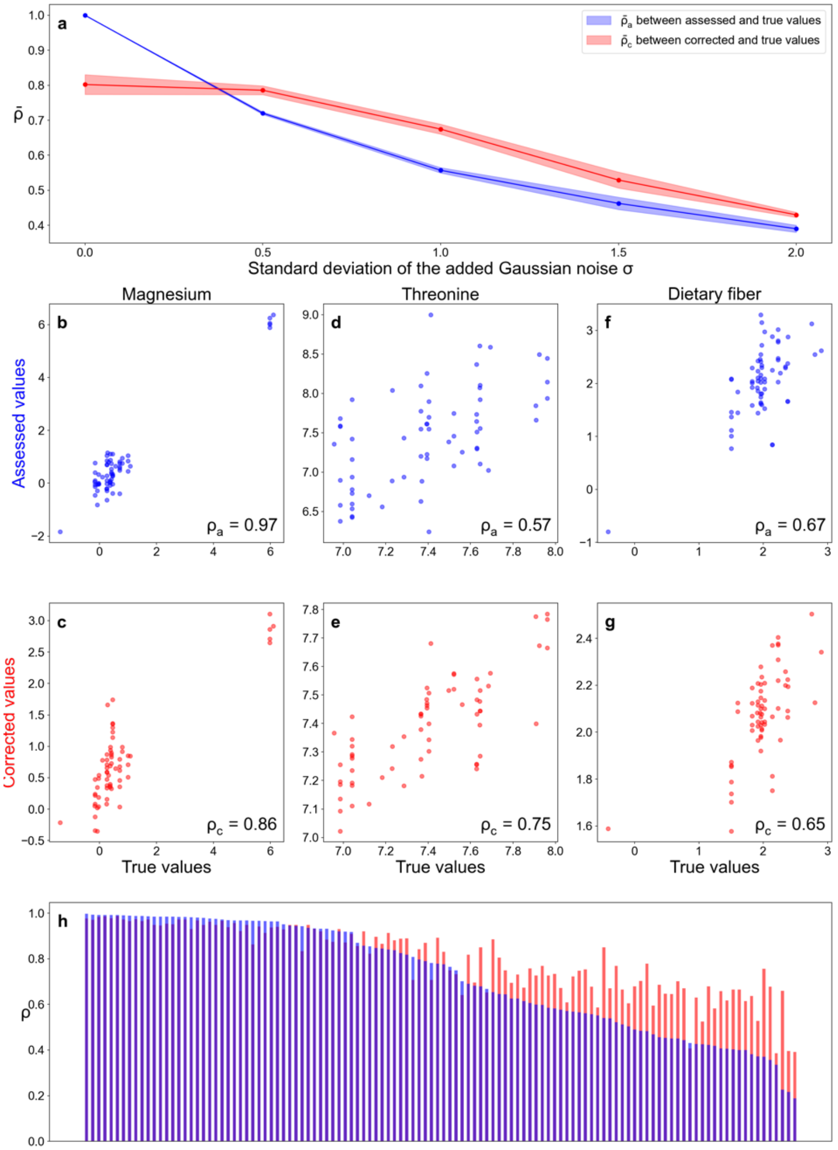


Supplementary Figure 5: **METRIC remains effective in correcting measurement errors in nutrients with large random errors, despite the overall poor correction performance when $\sigma = 0.5$ observed in the dataset WE-MACNUTR[39].** The Pearson's Rank Correlation Coefficient $\rho$ is adopted to evaluate the correlation across various types of nutrient profiles. **a,** $\rho_c$ (i.e., $\rho$ between corrected and true values) and $\rho_a$ (i.e., $\rho$ between assessed and true values) decrease as the standard deviation of added Gaussian noise $\sigma$ increases. All nutrient concentrations are in the unit of grams. All corrected/true values shown are the log of nutrient concentrations. All following panels focus on the case of $\sigma$=1.0. **b,** The correlation between assessed values and true values of log concentrations of magnesium among different samples. **c,** The correlation between corrected values (predictions of METRIC) and true values of log concentrations of magnesium among different samples. **d-e,** The similar comparison for threonine shows a modest correction. **f-g,** The similar comparison for dietary fiber shows no correction. **h,** The correction performance for all nutrients is measured by $(\rho_c - \rho_a)$. Nutrient names are not added due to lack of space.

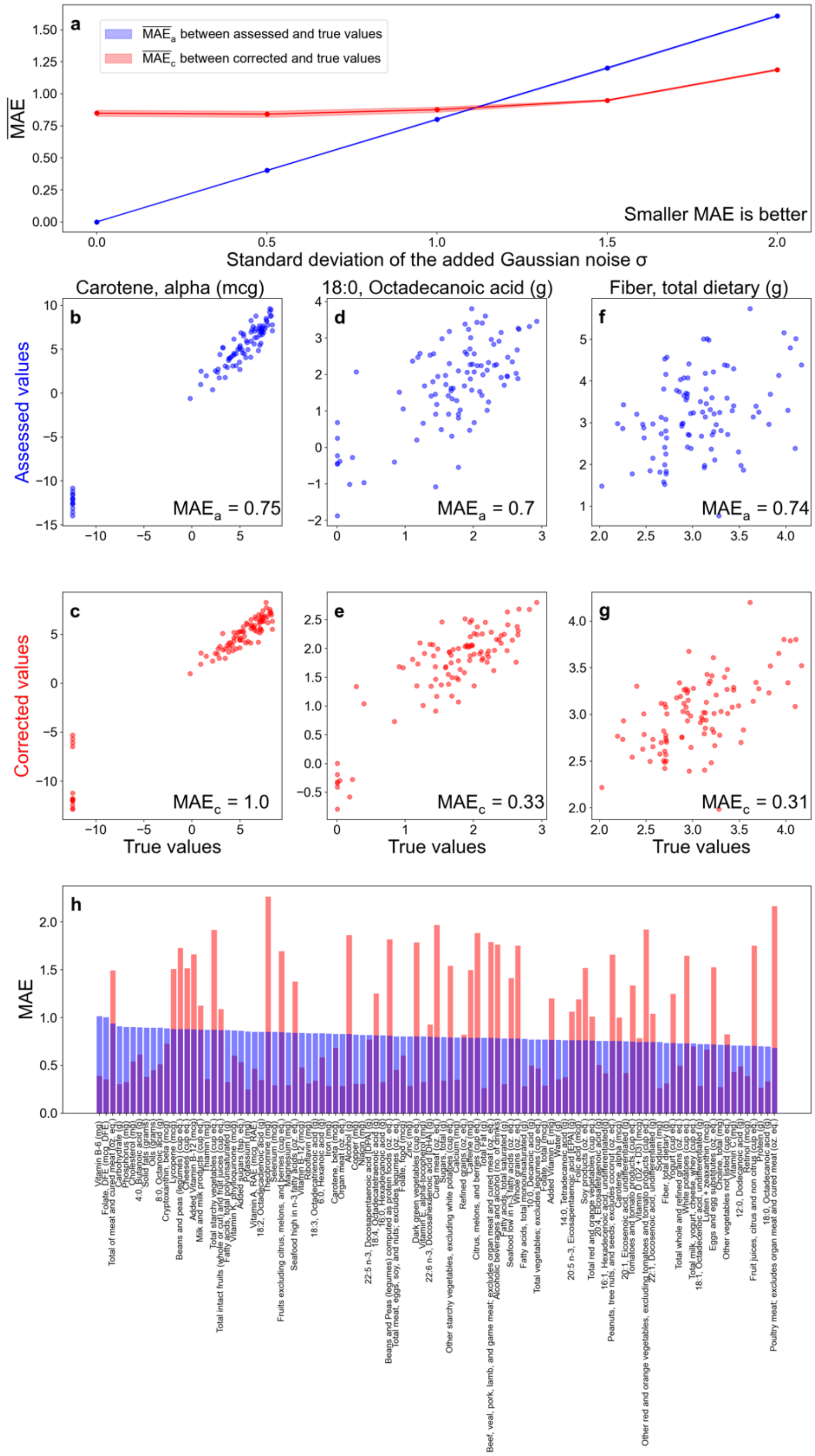


Supplementary Figure 6: **METRIC can correct the measurement error in assessed nutrient profiles on real data from MCTS[31].** The mean squared error (MAE) is adopted to evaluate the agreement between different types of nutrient profiles. All nutrient concentrations are in the unit of grams. All corrected/true values shown are the log of nutrient concentrations. **a,** $MAE_c$ (i.e., MAE between corrected and true values) and $MAE_a$ (i.e., MAE between assessed and true values) increase as the standard deviation of added Gaussian noise $\sigma$ increases. ($MAE_a -$

$MAE_c$) increases as $\mu$ increases. All following panels focus on the case of $\sigma$=1.0. **b,** The MAE between assessed values and true values of log concentrations of carotene among different samples. **c,** The MAE between corrected values (predictions of METRIC) and true values of log concentrations of carotene among different samples. **d-e,** The similar comparison for octadecanoic acid shows a good correction. **f-g,** The similar comparison for fiber shows a strong correction. **h,** The correction performance for all nutrients is measured by ($MAE_a - MAE_c$).

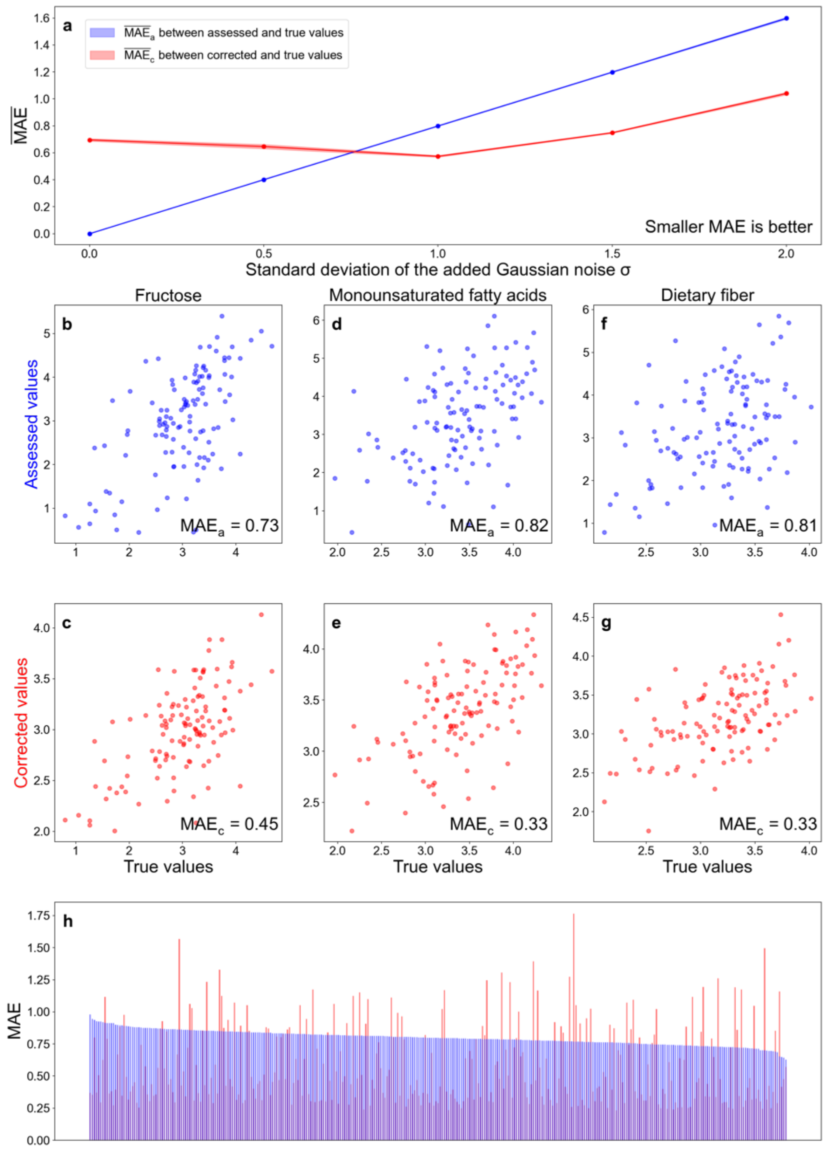


Supplementary Figure 7: **METRIC can correct the measurement error in assessed nutrient profiles from MLVS[36,37].** The mean squared error (MAE) is adopted to evaluate the agreement between different types of nutrient profiles. All nutrient concentrations are in the unit of grams. All corrected/true values shown are the log of nutrient concentrations. **a,** $MAE_c$ (i.e., MAE between corrected and true values) and $MAE_a$ (i.e., MAE between assessed and true values) increase as the standard deviation of added Gaussian noise $\sigma$ increases. $(MAE_a - MAE_c)$ increases as $\mu$ increases. All following panels focus on the case of $\sigma$=1.0. **b,** The MAE between assessed values and true values of log concentrations of fructose among different samples. **c,** The MAE between corrected values (predictions of METRIC) and true values of log concentrations of fructose among different samples. **d-e,** The similar comparison for monounsaturated fatty acids shows a great correction. **f-g,** The similar comparison for dietary fiber shows a strong correction. **h,** The correction performance for all nutrients is measured by $(MAE_a - MAE_c)$. Nutrient names are not added due to lack of space.

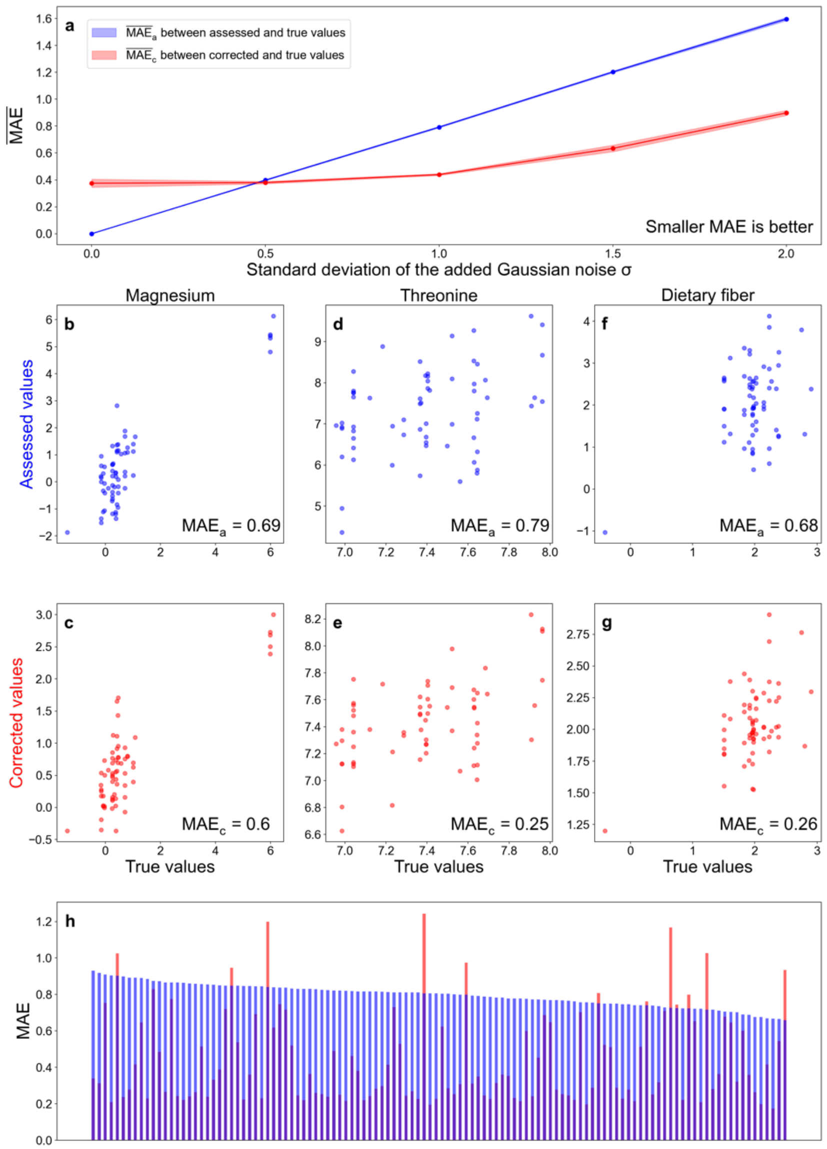


Supplementary Figure 8: **METRIC can correct the measurement error in assessed nutrient profiles from WE-MACNUTR[39].** The mean squared error (MAE) is adopted to evaluate the agreement between different types of nutrient profiles. All nutrient concentrations are in the unit of grams. All corrected/true values shown are the log of nutrient concentrations. **a,** $MAE_c$ (i.e., MAE between corrected and true values) and $MAE_a$ (i.e., MAE between assessed and true values) increase as the standard deviation of added Gaussian noise $\sigma$ increases. $(MAE_a - MAE_c)$ increases as $\mu$ increases. All following panels focus on the case of $\sigma$=1.0. **b,** The MAE between assessed values and true values of log concentrations of magnesium among different samples. **c,** The MAE between corrected values (predictions of METRIC) and true values of log concentrations of magnesium among different samples. **d-e,** The similar comparison for threonine shows a great correction. **f-g,** The similar comparison for dietary fiber shows a strong correction. **h,** The correction performance for all nutrients is measured by $(MAE_a - MAE_c)$. Nutrient names are not added due to lack of space.

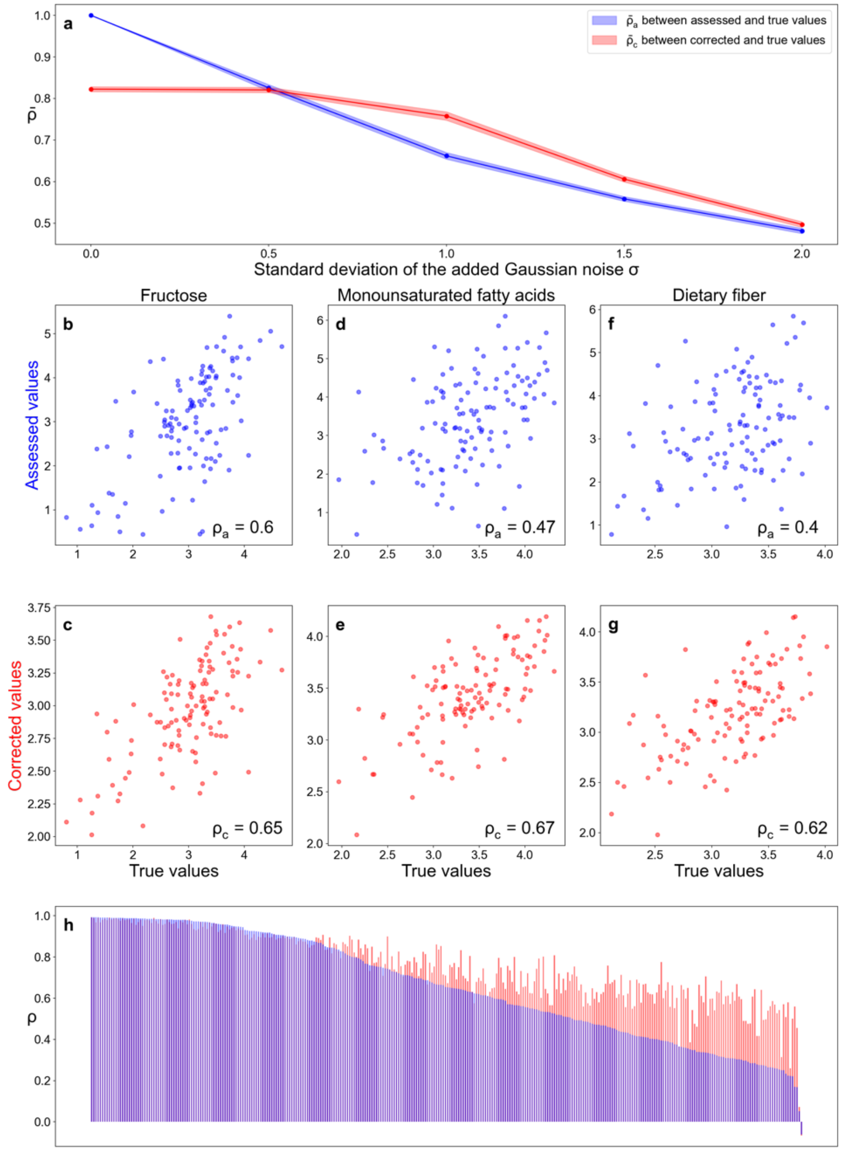


Supplementary Figure 9: **METRIC can correct the measurement error in assessed nutrient profiles from MLVS[36,37] without using gut microbial compositions in the input.** The Pearson's Rank Correlation Coefficient $\rho$ is adopted to evaluate the correlation across various types of nutrient profiles. All nutrient concentrations are in the unit of grams. All corrected/true values shown are the log of nutrient concentrations. **a,** $\rho_c$ (i.e., $\rho$ between corrected and true values) and $\rho_a$ (i.e., $\rho$ between assessed and true values) decrease as the standard deviation of added Gaussian noise $\sigma$ increases. All following panels focus on the case of $\sigma$=1.0. **b,** The correlation between assessed values and true values of log concentrations of fructose among different samples. **c,** The correlation between corrected values (predictions of METRIC) and true values of log concentrations of fructose among different samples. **d-e,** The similar comparison for monounsaturated fatty acids shows a modest correction. **f-g,** The similar comparison for dietary fiber shows a strong correction. **h,** The correction performance for all nutrients is measured by $(\rho_c - \rho_a)$. Nutrient names are not added due to lack of space.

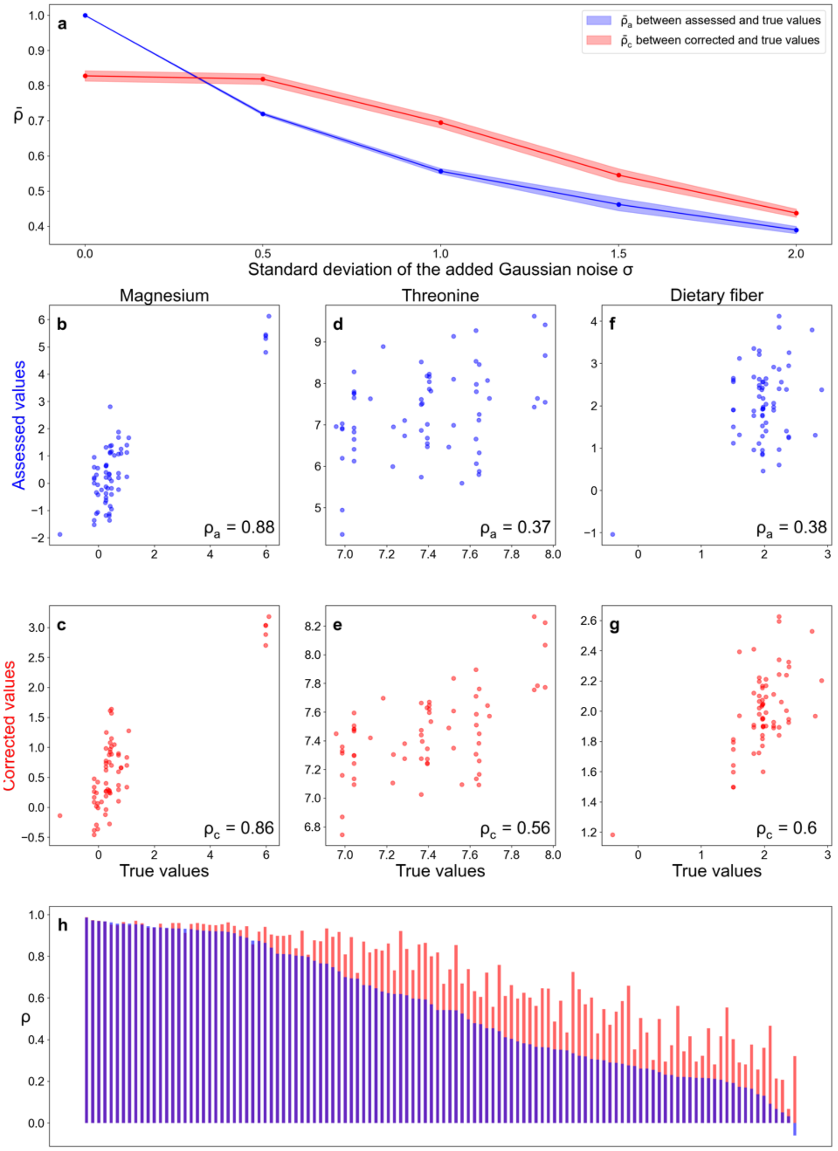


Supplementary Figure 10: **METRIC can correct the measurement error in assessed nutrient profiles from WE-MACNUTR[39] without using gut microbial compositions in the input.** The Pearson's Rank Correlation Coefficient $\rho$ is adopted to evaluate the correlation across various types of nutrient profiles. **a,** $\rho_c$ (i.e., $\rho$ between corrected and true values) and $\rho_a$ (i.e., $\rho$ between assessed and true values) decrease as the standard deviation of added Gaussian noise $\sigma$ increases. All nutrient concentrations are in the unit of grams. All corrected/true values shown are the log of nutrient concentrations. All following panels focus on the case of $\sigma$=1.0. **b,** The correlation between assessed values and true values of log concentrations of magnesium among different samples. **c,** The correlation between corrected values (predictions of METRIC) and true values of log concentrations of magnesium among different samples. **d-e,** The similar comparison for threonine shows a modest correction. **f-g,** The similar comparison for dietary fiber shows a strong correction. **h,** The correction performance for all nutrients is measured by $(\rho_c - \rho_a)$. Nutrient names are not added due to lack of space.

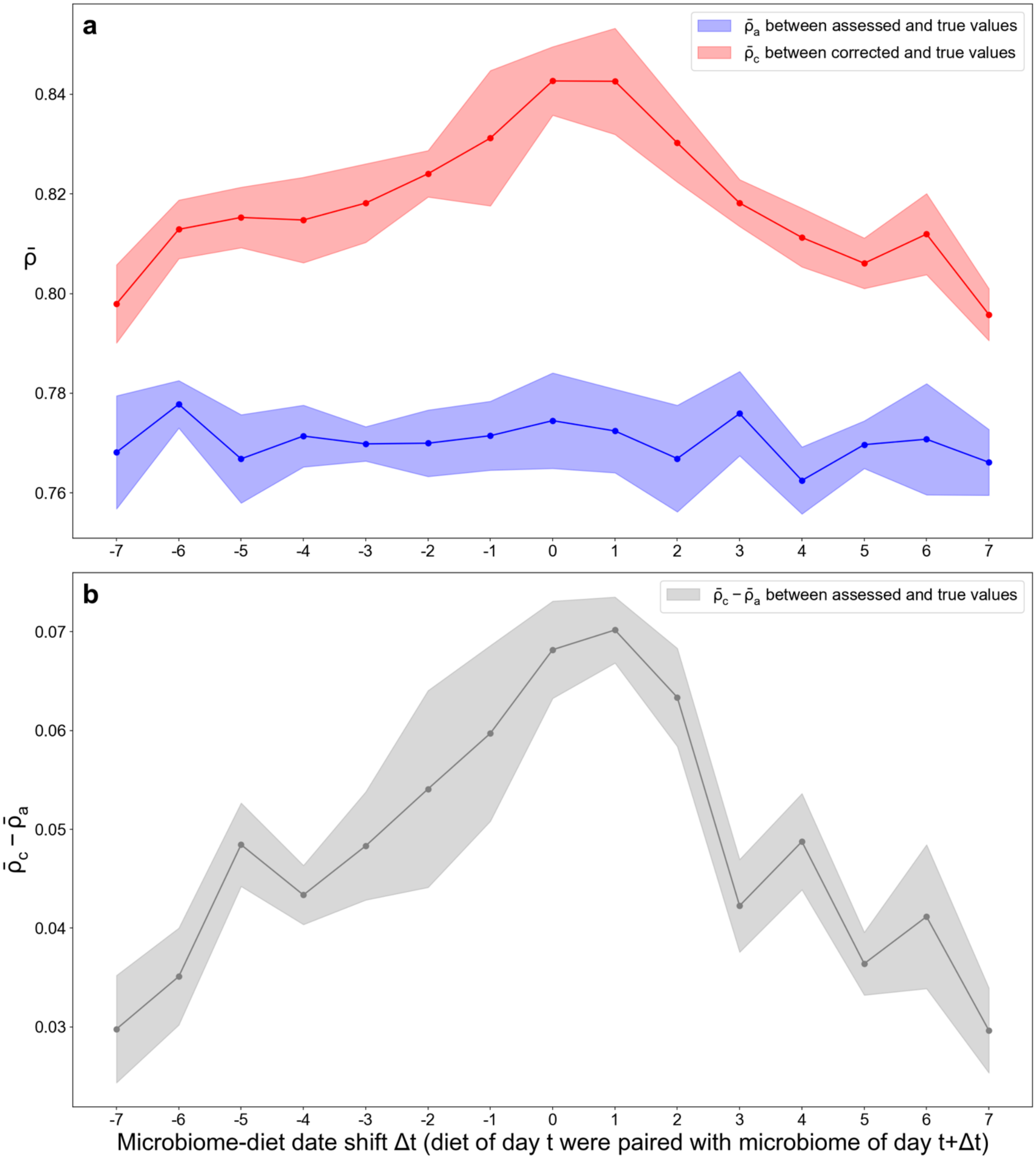


Supplementary Figure 11: **METRIC remains most effective when the microbiome-diet date shift is small for the dataset MCTS[31].** The microbiome-diet date shift $\Delta t$ is achieved by pairing the diet of day $t$ with the microbiome of day $t + \Delta t$. All following panels focus on the case of $\sigma$=1.0. The Pearson's Rank Correlation Coefficient $\rho$ is adopted to evaluate the correlation across various types of nutrient profiles. **a,** $\rho_c$ (i.e., $\rho$ between corrected and true values) changes and $\rho_a$ (i.e., $\rho$ between assessed and true values) does not change much as the microbiome-diet date shift $\Delta t$ changes. **b,** The correction performance for all nutrients ($\rho_c - \rho_a$) is strong when the microbiome-diet date shift $\Delta t$ is around 1 day.

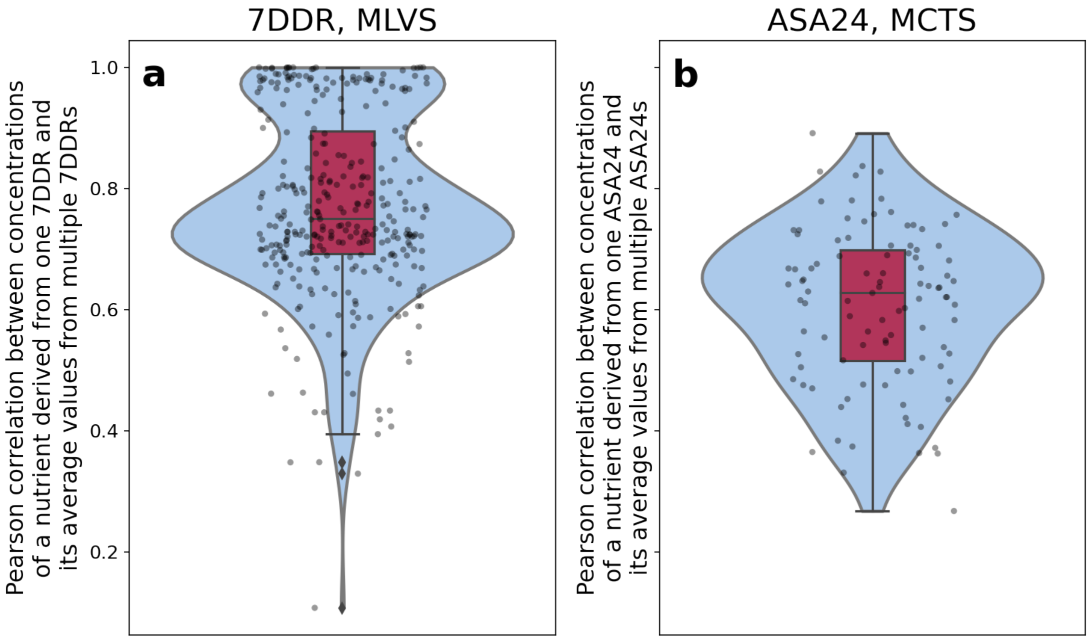


Supplementary Figure 12: **The distribution of Pearson's rank correlation coefficients between nutrient concentrations from a single-day dietary assessment and average values from multiple-day dietary assessments. a,** The distribution of Pearson's rank correlation coefficients between nutrient concentrations from a single-day 7DDR and average values from multiple-day 7DDRs in the dataset MLVS[36,37]. **b,** The distribution of Pearson's rank correlation coefficients between nutrient concentrations from a single-day ASA24 and average values from multiple-day ASA24 in the dataset MCTS[31]**.** In the boxplot, the black line in the middle of the red box is the median, the lower and upper hinges correspond to the first and third quartiles, and the black line ranges from the 1.5 × IQR (where IQR is the interquartile range) below the lower hinge to 1.5 × IQR above the upper hinge. The violin plot is smoothed by a kernel density estimator, with 0 and 1 set as the lower and upper bound, respectively. The Pearson's rank correlation coefficient for each nutrient is shown as a black dot.

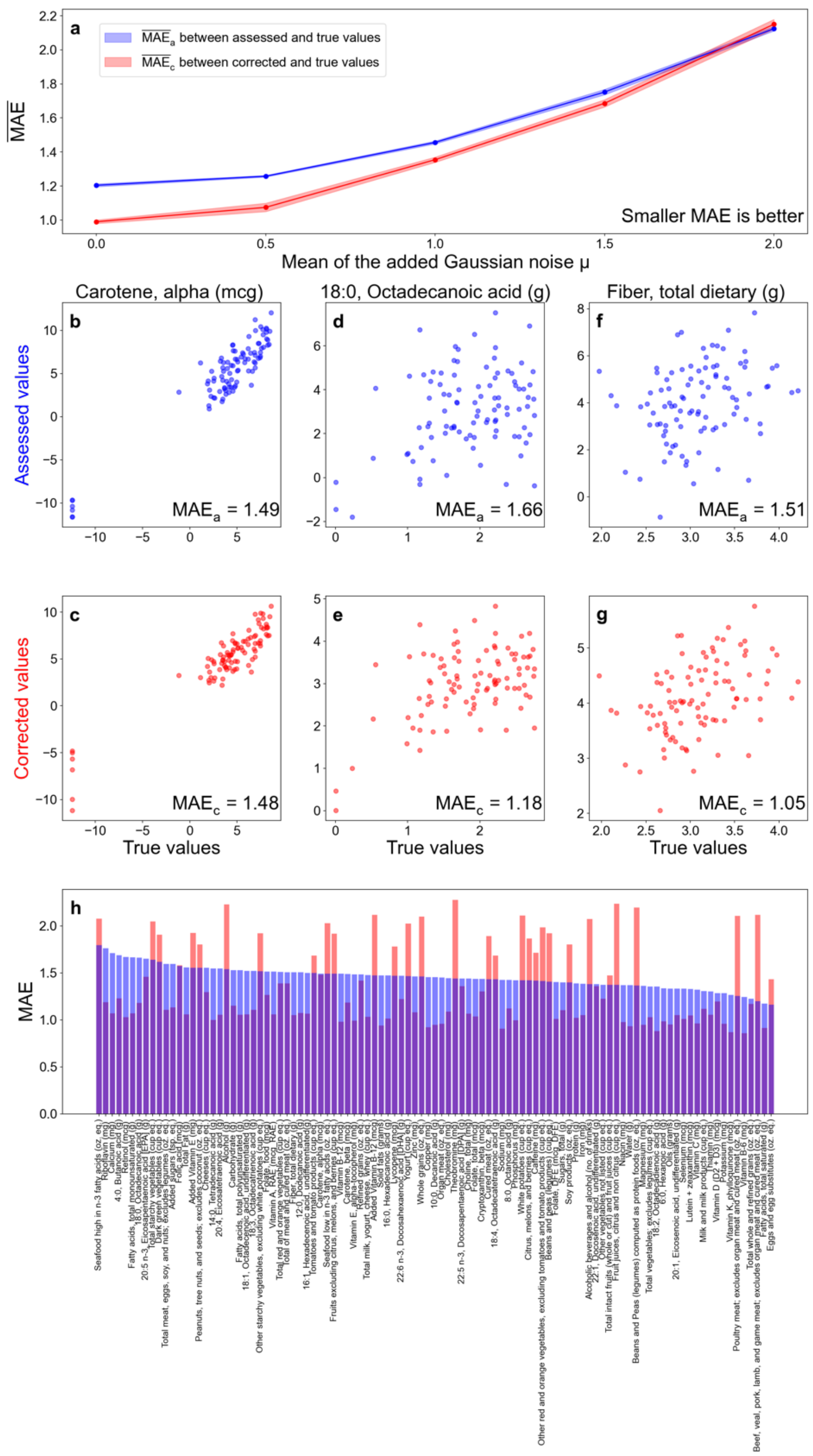


Supplementary Figure 13: **METRIC can correct the random part of measurement error with a nonzero mean in assessed nutrient profiles on real data from MCTS[31].** The standard deviation of added Gaussian noise $\sigma$ is set as 1, while the mean of added Gaussian noise $\mu$ varies. The mean squared error (MAE) is adopted to evaluate the agreement between different types of nutrient profiles. All nutrient concentrations are in the unit of grams. All corrected/true values shown are the log of nutrient concentrations. **a,** $MAE_c$ (i.e., MAE between corrected and true values) and $MAE_a$ (i.e., MAE between assessed and true values) increase as the mean of added Gaussian noise $\mu$ increases. $(MAE_a - MAE_c)$ shrinks to zero as $\mu$ increases. All following panels focus on the case of $\mu$=1.0. **b,** The MAE between assessed values and true values of log concentrations of carotene among different samples. **c,** The MAE between corrected values (predictions of METRIC) and true values of log concentrations of carotene among different samples. **d-e,** The similar comparison for octadecanoic acid shows a great correction. **f-g,** The similar comparison for fiber shows a strong correction. **h,** The correction performance for all nutrients is measured by $(MAE_a - MAE_c)$.

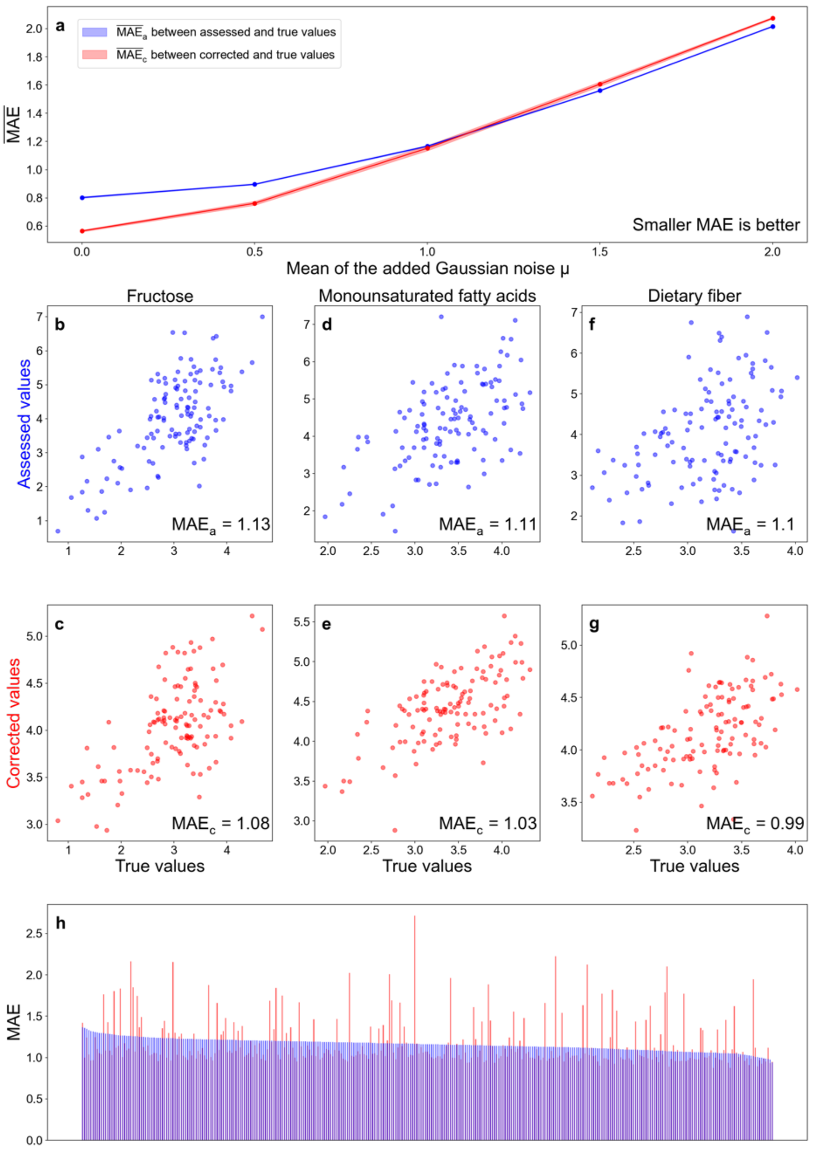


Supplementary Figure 14: **METRIC can correct the random part of measurement error with a nonzero mean in assessed nutrient profiles on real data from MLVS[36,37].** The standard deviation of added Gaussian noise $\sigma$ is set as 1, while the mean of added Gaussian noise $\mu$ varies. The mean squared error (MAE) is adopted to evaluate the agreement between different types of nutrient profiles. All nutrient concentrations are in the unit of grams. All corrected/true values shown are the log of nutrient concentrations. **a,** $MAE_c$ (i.e., MAE between corrected and true values) and $MAE_a$ (i.e., MAE between assessed and true values) increase as the mean of added Gaussian noise $\mu$ increases. All following panels focus on the case of $\mu$=1.0. $(MAE_a - MAE_c)$ shrinks to zero as $\mu$ increases. **b,** The MAE between assessed values and true values of log concentrations of fructose among different samples. **c,** The MAE between corrected values (predictions of METRIC) and true values of log concentrations of fructose among different samples. **d-e,** The similar comparison for monounsaturated fatty acids shows a great correction. **f-g,** The similar comparison for dietary fiber shows a strong correction. **h,** The correction performance for all nutrients is measured by $(MAE_a - MAE_c)$. Nutrient names are not added due to lack of space.

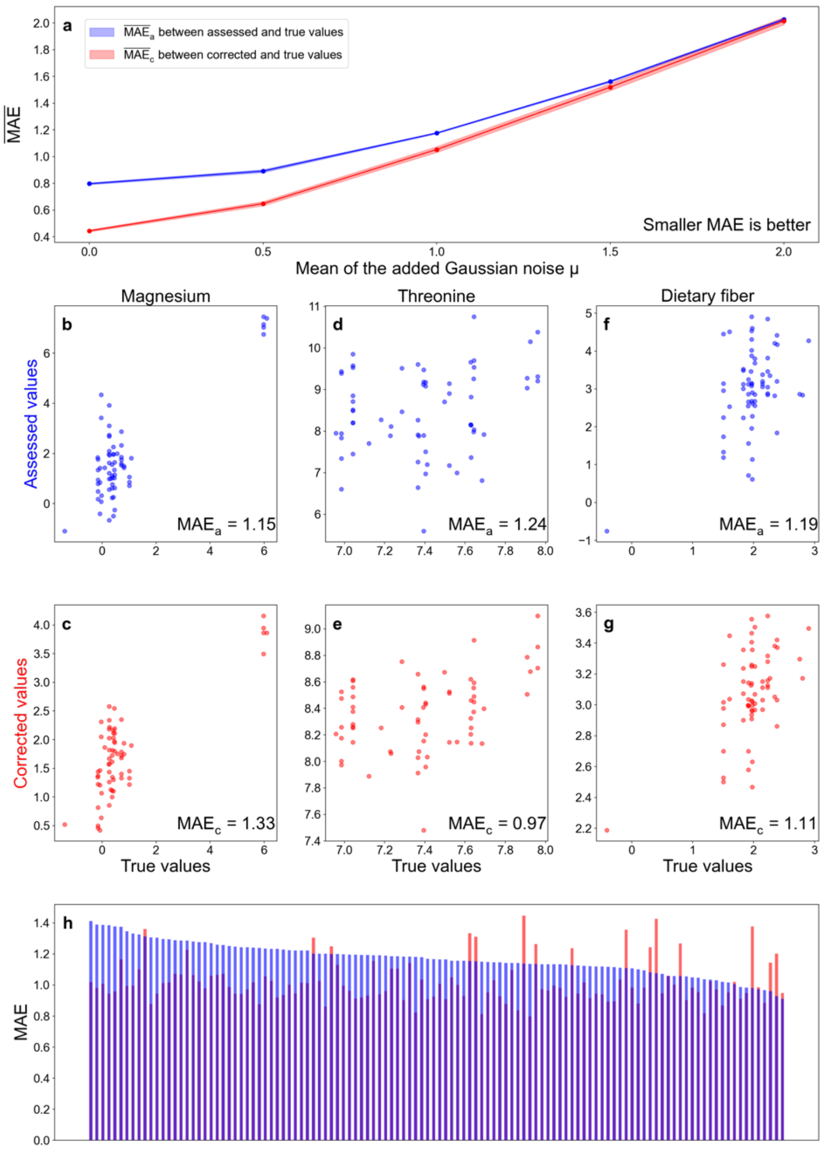


Supplementary Figure 15: **METRIC can correct the random part of measurement error with a nonzero mean in assessed nutrient profiles on real data from WE-MACNUTR[39].** The standard deviation of added Gaussian noise $\sigma$ is set as 1, while the mean of added Gaussian noise $\mu$ varies. The mean squared error (MAE) is adopted to evaluate the agreement between different types of nutrient profiles. All nutrient concentrations are in the unit of grams. All corrected/true values shown are the log of nutrient concentrations. **a,** $MAE_c$ (i.e., MAE between corrected and true values) and $MAE_a$ (i.e., MAE between assessed and true values) increase as the mean of added Gaussian noise $\mu$ increases. $(MAE_a - MAE_c)$ shrinks to zero as $\mu$ increases. All following panels focus on the case of $\mu$=1.0. **b,** The MAE between assessed values and true values of log concentrations of magnesium among different samples. **c,** The MAE between corrected values (predictions of METRIC) and true values of log concentrations of magnesium among different samples. **d-e,** The similar comparison for threonine shows a great correction. **f-g,** The similar comparison for dietary fiber shows a strong correction. **h,** The correction performance for all nutrients is measured by $(MAE_a - MAE_c)$. Nutrient names are not added due to lack of space.

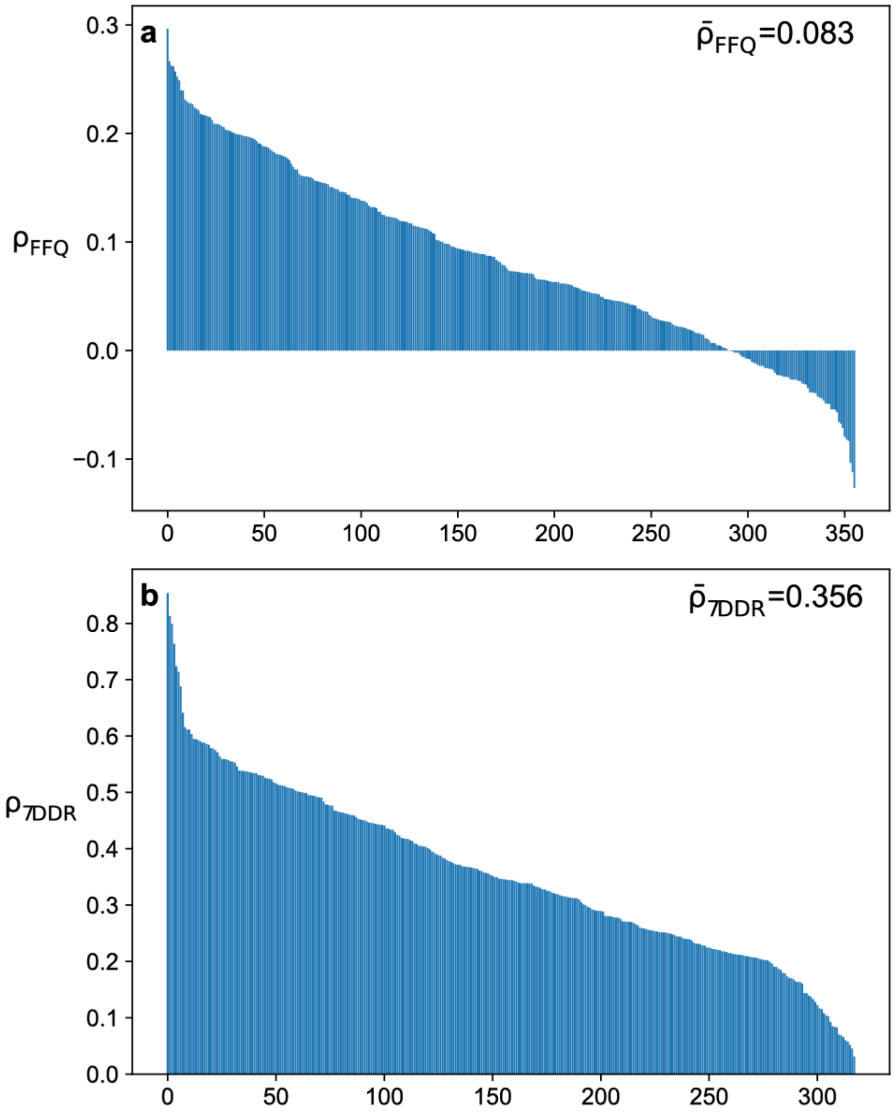


Supplementary Figure 16: **Using gut microbial compositions as the input, the predictability of FFQ-based nutrient profiles is much worse than that of 7DDR-based nutrient profiles for MLVS[36,37].** For both prediction tasks, a multilayer perceptron model with 3 hidden layers, each with 256 nodes (the same as METRIC) is used. To assess the predictability of nutrient profiles, Pearson's Rank Correlation Coefficient $\rho$ between the true and predicted nutrient concentrations is computed. **a,** The rank plot of Pearson's Rank Correlation Coefficient $\rho_{FFQ}$ between the true and predicted nutrient concentrations derived from FFQ. The mean value $\bar{\rho}_{FFQ} = 0.083$. **b,** The rank plot of Pearson's Rank Correlation Coefficient $\rho_{7DDR}$ between the true and predicted nutrient concentrations derived from 7DDR. The mean value $\bar{\rho}_{7DDR} = 0.356$.

#### Supplementary Data Legends

**Supplementary Data 1:** The inferred sensitivity values for the dataset MCTS.
